# Exploring evolution to uncover insights into protein mutational stability

**DOI:** 10.1101/2024.05.28.596203

**Authors:** Pauline Hermans, Matsvei Tsishyn, Martin Schwersensky, Marianne Rooman, Fabrizio Pucci

## Abstract

Determining the impact of mutations on the thermodynamic stability of proteins is essential for a wide range of applications such as rational protein design and genetic variant interpretation. Since protein stability is a major driver of evolution, evolutionary data are often used to guide stability predictions. Many state-of-the-art stability predictors extract evolutionary information from multiple sequence alignments (MSA) of proteins homologous to a query protein, and leverage it to predict the effects of mutations on protein stability. To evaluate the power and the limitations of such methods, we used the massive amount of stability data recently obtained by deep mutational scanning to study how best to construct MSAs and optimally extract evolutionary information from them. We tested different evolutionary models and found that, unexpectedly, independent-site models achieve similar accuracy to more complex epistatic models. A detailed analysis of the latter models suggests that their inference often results in noisy couplings, which do not appear to add predictive power over the independent-site contribution, at least in the context of stability prediction. Interestingly, by combining any of the evolutionary features with a simple structural feature, the relative solvent accessibility of the mutated residue, we achieved similar prediction accuracy to supervised, machine learning-based, protein stability change predictors. Our results provide new insights into the relationship between protein evolution and stability, and show how evolutionary information can be exploited to improve the performance of mutational stability prediction.

## 1 Introduction

Understanding how mutations impact protein thermodynamic stability is of fundamental interest for a wide range of applications spanning from protein design [1, 2] to the interpretation of genetic variants [3, 4]. Evolution has largely been used to guide computational or experimental mutagenesis aimed at improving protein stability, the latter being the primary driver of evolution [5, 6, 7].

Directed evolution methods, which aim to replicate evolutionary processes *in vitro* through iterative steps of random mutagenesis, selection and amplification, have played a fundamental role in the (re)design of new proteins with increased stability [8, 9, 10]. Computational mutagenesis approaches have been extensively improved in the last decade [2, 11, 12, 13, 14] and have often been combined with experimental methods to limit the time-consuming exploration of the vast protein sequence space [1, 15]. Recently, deep learning techniques have also been introduced to the field, but their application has often been constrained by the limited availability of experimental thermodynamic data. New high-throughput stability assays, such as the assay recently developed in [16], are currently changing the field by generating folding stability measurements on an impressively large scale, providing essential information to better understand protein stability and to constitute sufficiently large datasets to train deep learning models.

Some current state-of-the-art models for stability prediction [17, 18, 19] already take advantage of evolution and of the huge amount of sequence data available in metagenomic databases [20]. They extract evolutionary information from multiple sequence alignments (MSAs) of proteins homologous to a query protein, which is used to predict how mutations affect protein stability. Although evolution and protein stability are known to be closely related [21], they are not always directly linked. Indeed, functional regions such as catalytic or binding sites are not at all optimized for stability, whereas they clearly show a very strong conservation [22, 23]. In addition, several open questions are still being debated in the field. For example, it remains unclear whether the effects of mutations are conserved throughout evolution [24, 25]. This question is related to the intriguing role of epistatic effects [25] in shaping protein evolution and stability, which is still not fully understood. While some evidence points to their essential role [26, 27, 28], other studies suggest otherwise [24]. From a computational perspective, the inclusion of epistatic contributions in evolutionary models generally seems to better capture the protein mutational fitness landscape [29, 30], though this is not always the case and often depends on the specific protein considered [31].

Finally, from a practical point of view, it is unclear what is the best way to generate and manage MSAs in order to extract evolutionary signals that can improve protein stability prediction and protein design. In the last couple of years, much attention has been paid to optimizing or sub-sampling the input MSAs to improve downstream tasks such as predicting the three-dimensional (3D) structures of proteins and of their complexes [32, 33, 34, 35, 36]. A recent analysis on how MSA construction impacts the outcome of mutational fitness prediction [37] presented interesting findings, in particular that prediction accuracy does not always correlate with alignment depth; evolutionary information extracted from shallow MSAs can sometimes predict the effect of mutations on protein fitness with high accuracy. Adding homologous sequences that are evolutionary distant from the query sequence does thus not necessarily improve predictions [37, 38].

We leveraged the massive amount of protein stability data generated in [16] and analyzed how to optimally extract evolutionary information from MSAs for mutational stability predictions. One of our goals being to gain insight into the relationship between protein stability and evolution, we compared the ability of various kinds of evolutionary models to predict protein stability changes upon mutations. We also highlighted the relationships between residue solvent accessibility (RSA) which is a simple structural feature, the evolutionary conservation of these residues, and the effect of their substitutions on protein stability. Based on these relationships, we devised a way to modulate evolutionary features with RSA, resulting in simple unsupervised models that yield results similar to those of more complex state-of-the-art protein stability change predictors.

## 2 Methods

### 2.1 Multiple Sequence Alignment Construction

For each query sequence, we generated MSAs using the iterative JackHMMER method [39] against a given sequence dataset. Basically, JackHMMER performs a quick sequence similarity search on the considered sequence dataset to build a first MSA, which is used to set up a hidden Markov model (HMM). This HMM is then searched against the sequence dataset to select new sequences and construct the next MSA and HMM, and so on. Each iteration refines both the MSA and the HMM used for its construction.

In order to assess the relevance of the MSAs for stability change predictions and compare different ways of constructing them, we tested various numbers of iterations and E-value thresholds in JackHMMER (see Section 3.1). We also evaluated different sequence datasets. We primarily used the UniRef90 set [40], but also tested searching in two larger datasets: UniRef100 and Metagenomics, a metagenomic dataset constructed by merging UniRef90 [40], the Big Fantastic Database [41], the Gut Phage Database [42], MetaEuk [43], the Metagenomic Gut Virus (MGV) [44], SMAG [45], TOPAZ [46] and MGnify [47].

As MSAs can contain clusters of very closely related proteins, measuring the amount of information contained in them by *N*_tot_, the total number of sequences it contains, can sometimes be inaccurate. For that reason, we rather used *N*_eff_, the effective number of independent sequences, defined as the sum of the weights *w*_*j*_ assigned to each sequence *s*_*j*_ in the MSA [48]:

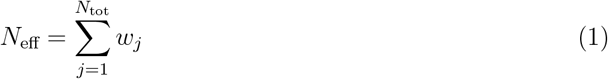

in order to assess the impact of the “depth” of an MSA on predictions. The weight *w*_*j*_ of a sequence *s*_*j*_ is defined as the inverse of *m*_*j*_, the number of sequences in the MSA sharing at least 80% identity with *s*_*j*_:

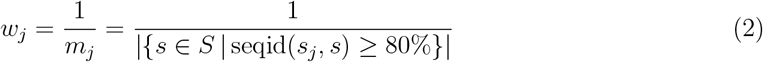

where *S* is the set of sequences in the MSA. In this way, sequences without similar sequences in the dataset have a weight of one, and sequences with similar sequences are down-weighted. These weights were computed as in [48] using the **plmc** software from the EVcouplings suite [30].

### 2.2 Variant Dataset Construction

We used the recent deep mutational scanning dataset constructed in [16], which contains about one million experimental estimations of protein folding free energy (Δ*G*) obtained through cDNA display proteolysis applied to 396 “wild-type” sequences. We first curated this dataset by averaging duplicate measured Δ*G* values for specific sequences (both “wild-type” and “mutant”). We focused on single amino acid substitutions and computed the corresponding changes in folding free energy upon mutations, defined as ΔΔ*G* = Δ*G*_*mt*_ − Δ*G*_*wt*_, where *wt* and *mt* stand for “wild-type” and for “mutant” respectively.

To reduce protein-specific imbalances in the dataset, we employed the clustering method of CD-HIT [49] with default parameters (sequence identity threshold of 0.9). In this way, we removed closely related sequences and limited the number of “wild-type” sequences to 308. As many of these proteins have a very small number of known homologs (among which *de novo* designed proteins), we further refined the dataset by removing proteins whose MSA (obtained by JackHMMER on UniRef90 with one iteration and default parameters) contains less than 100 sequences. The resulting dataset, called 𝒟, contains 135, 056 ΔΔ*G* values of single-site mutations in 129 protein domains with lengths ranging from 32 to 72 amino acids. The obtained distribution of the experimental ΔΔ*G* values is presented in Supplementary Section 1. As expected, most of the mutations are destabilizing, with an average destabilizing ΔΔ*G* of 0.82 kcal/mol.

In order to investigate the effect of MSA depth on different evolutionary models, we used the *N*_eff_ value (Eq. 1) as a criterion to partition the dataset 𝒟 into subsets: 𝒟_10_ (10 ≤ *N*_eff_ *<* 100, 17 proteins), 𝒟_100_ (100 ≤ *N*_eff_ *<* 1000, 27 proteins), 𝒟_1000_ (1000 ≤ *N*_eff_ *<* 10000, 41 proteins) and 𝒟_10000_, (10000 ≤ *N*_eff_, 44 proteins). Values of *N*_eff_ were derived on MSAs constructed on UniRef90 with two JackHMMER iterations and an E-value threshold of 10^−7^, as we showed in Section 3.1 that these parameter values are optimal for stability predictions. The structures of all “wild-type” proteins were modeled using AlphaFold2 [33].

To analyze the generalizability of our results to larger proteins, we additionally used the recently published literature-based, non-systematic, dataset S4038 [50]. We constructed dataset ℒ by selecting the 7 proteins in S4038 which are larger than the proteins from 𝒟 and for which experimental ΔΔ*G* values of at least 60 mutations have been reported (see Supplementary Section 2 for details).

The complete datasets as well as the PDB structures used in this paper are available in our GitHub repository https://github.com/3BioCompBio/EvoStability.

### 2.3 Residue and Mutation Properties

We first defined three evolution-based features derived from positional amino acid frequencies in an MSA. To avoid divergent values and manage the lack of information in small MSAs, we defined a way to regularize the observed amino acid frequencies. Considering the positional frequency *f*_*i*_(*a*) of amino acid *a* at position *i* in the MSA and the frequency *f* (*a*) of amino acid *a* in the full MSA, we defined the regularized frequencies 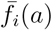 and 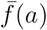 as:

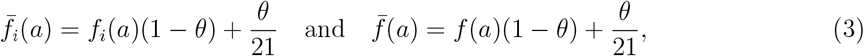

where *θ* is a regularization factor that we set to 0.01, and 21 stands for the number of possible states (20 standard amino acids, and 1 gap). Using these regularized frequencies, the three evolution-based features are:

- **Conservation Index (CI)** [51]. It is a measure of the conservation of residues at each position in an MSA:

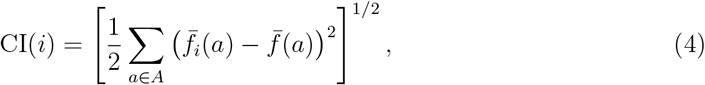

where *A* is the set of 20 amino acids.
- **Log-Odd Ratio (LOR)** [52]. By comparing the frequencies of wild-type (*wt*) and mutant (*mt*) amino acids in the MSA at a given position, we can statistically estimate their prevalence in their evolutionary context. The log-odd ratio of observing the *wt* amino acid with respect to the *mt* amino acid at position *i* is given by:

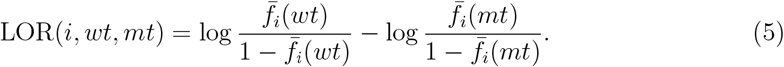

The sign of LOR is set such that a mutation from a highly represented amino acid *wt* to a poorly represented amino acid *mt* has a positive LOR, which is comparable to a destabilizing mutation with ΔΔ*G >* 0.
- **Weighted Log-Odd Ratio (LOR**_*w*_). To reduce the impact of clusters of closely related sequences in the MSA, we slightly modified the definition of LOR by rectifying the count for amino acid frequencies using the weights assigned to each sequence in the MSA (computed with **plmc** as for *N*_eff_, see Section 2.1). The weighted log-odd ratio is defined as:

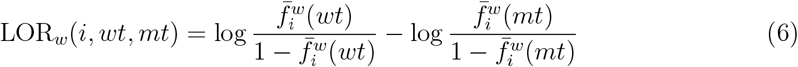

where the weighted frequency 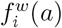 of an amino acid *a* at position *i* is defined as:

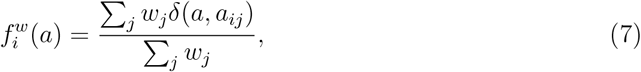

with *w*_*j*_ is the weight of sequence *j, a*_*ij*_ the amino acid at position *i* in sequence *j* and *δ*(*A, B*) is the Kronecker symbol, which equals one if the *A* and *B* coincide, and zero otherwise. Note that in this case, the regularization step of Eq. 3 is applied to the weighted frequencies.

We also used four mutational coevolutionary methods:

- **pycofitness** [53] is a direct coupling analysis (DCA) based model. DCA provide a statistical representation of a family of homologous protein sequences from a given MSA. Let *S* = (*a*_1_, *a*_2_, …, *a*_*L*_) be a protein sequence of length *L*, where *a*_*i*_ is the amino acid type at position *i*. The sequence *S* is assumed to be sampled across evolution with a probability given by the Boltzmann distribution, *P* (*S*), which can be expressed as:

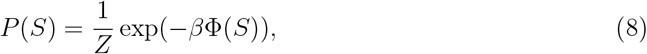

where *β* is the inverse temperature, *Z* denotes the partition function, and Φ(*S*) is the energy of the system:

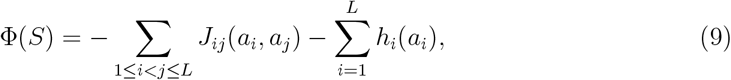

with *h*_*i*_(*a*_*i*_) and *J*_*ij*_(*a*_*i*_, *a*_*j*_) corresponding to single-site fields and coupling parameters, respectively. To infer them, pycofitness uses a pseudo-likelihood maximization approach [54] (see Supplementary Section 6 for details). Once the parameters are inferred, the effect of mutation of residue *a* into *b* at position *i* in the sequence, Δ*X*(*i, a, b*), is computed as :

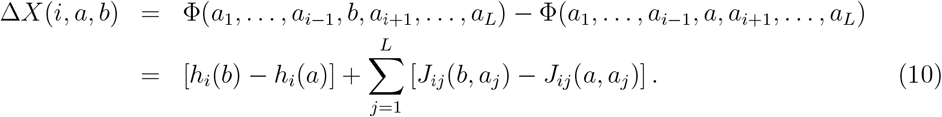
- **EVcouplings** [30, 55] suite contains two models for predicting the effects of mutations, the epistatic and the independent-site models, which are referred to as EVcouplings-epi and EVcouplings-ind, respectively. The EVcouplings-epi model accounts for epistatis by modeling interactions between all pairs of residues using a DCA model, with coefficients inferred from the input MSA through a pseudo-likelihood maximization approach [54]. This method is analogous to pycofitness. EVcouplings-ind is a site-wise maximum entropy model, which does not take into account interactions between sites.
- **ArDCA** [56] is an epistatic method that uses an autoregressive model to compute the full probability *P* (*S*) of observing the sequence *S* = (*a*_1_, …, *a*_*L*_) across evolution as the product of conditional probabilities *P* (*S*) = *P* (*a*_1_)*P* (*a*_2_|*a*_1_) … *P* (*a*_*L*_|*a*_*L*−1_, …, *a*_1_). The model parameters are inferred through a maximum-likelihood approach.
- **GEMME** [31] combines an epistatic and an independent-site version, which will be considered independently here. The epistatic model, referred to as GEMME-epi, infers the epistatic contributions from the MSA based on the minimal evolutionary distance between a sequence carrying the mutations and the query sequence in the evolutionary tree. The independent-site model, GEMME-ind, is based on the relative per-site frequencies of the wild-type and mutant residues.

Finally, we used one feature based on the protein 3D structure, and defined a way of combining it with the evolutionary features described above:

- **Relative Solvent Accessibility (RSA)**. The RSA of a residue at position *i*, RSA(*i*), is defined as the ratio (in %) of its solvent-accessible surface area in its 3D structure and in a Gly-X-Gly tripeptide in extended conformation [57]. We calculated it using our in-house software MuSiC [58], which uses an extension of the DSSP algorithm [59] and is available on the www.dezyme.com website.
- **RSA and evolution**

As mutations in the protein core tend to be more destabilizing than mutations at the protein surface (see Section 3.5), RSA is a crucial feature in estimating protein stability. For this reason, we devised a simple way of combining any mutational evolutionary score with the RSA as:

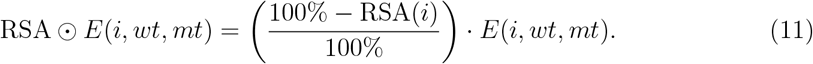

where *E*(*i, wt, mt*) is the evolutionary score of the substitution from residue *wt* to residue *mt* at position *i*.

## 3 Results

### 3.1 Impact of MSA construction and curation

We started by evaluating the impact of MSA construction and curation on the ability of independent-site evolutionary scores (i.e., LOR and LOR_*w*_, defined in Section 2.3) to predict changes in protein stability. In particular, we tested different protein sequence datasets, numbers of JackHMMER iterations, E-value thresholds and MSA curation criteria. As evaluation metric, we computed the average per-protein Spearman correlation coefficients between the evolutionary scores and the experimental ΔΔ*G* values from the variant dataset *D*. We present here the results obtained with LOR_*w*_ scores, as they turn out to be slightly more predictive than LOR values.

For the number of JackHMMER iterations (ranging from 1 to 7), corresponding to successive levels of MSA refinement, we found that MSAs obtained with 2 iterations yield LOR_*w*_ scores that correlate best with experimental ΔΔ*G* values (Fig. 1a). This is true regardless of the sequence dataset used. This result is statistically significant; the *p*-values for the comparison of correlations and the method to derive them are provided in Supplementary Section 3. Sequences added during the second iteration contribute to the diversity of the MSAs, which is beneficial for predicting changes in protein stability. Subsequent iterations appear to introduce noise in the MSA rather than adding any useful information.

**Figure 1:**
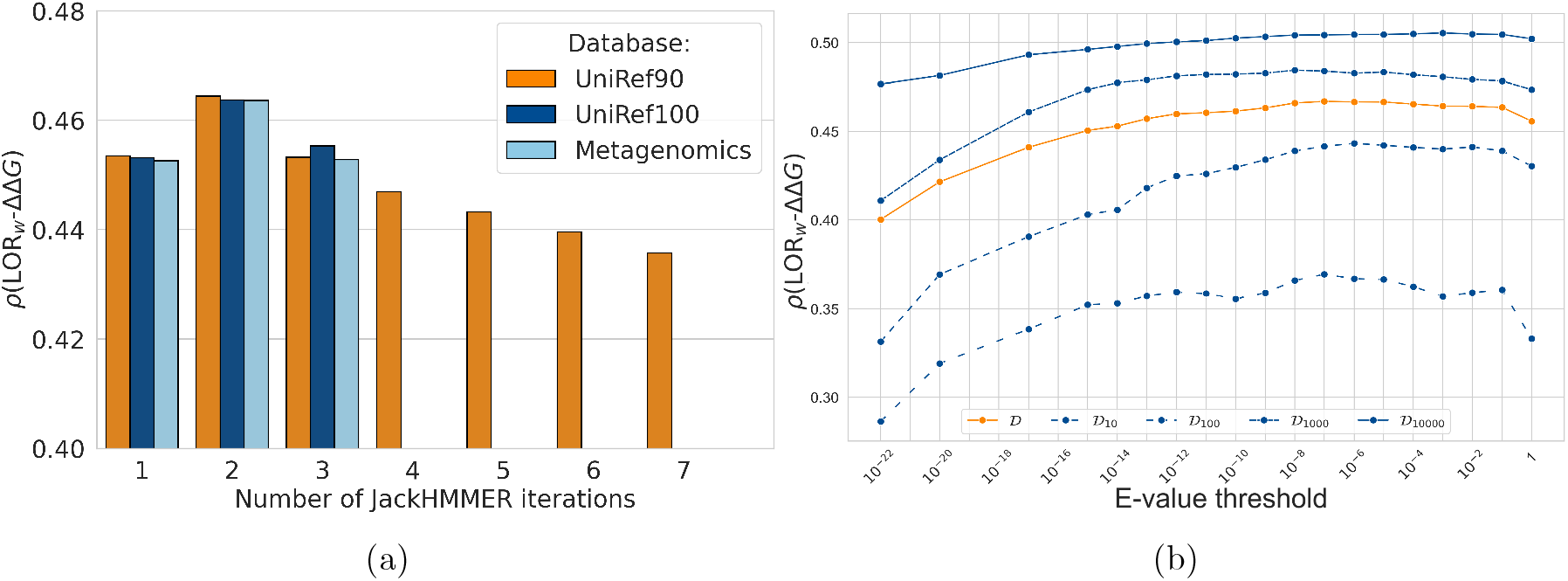
Average per-protein Spearman correlation coefficients *ρ* between LOR_*w*_ and ΔΔ*G* on dataset 𝒟 as a function of: (a) the protein sequence dataset and the number of JackHMMER iterations and (b) the E-value thresholds between 10^−22^ and 1 using UniRef90 and two JackHMMER iterations (on 𝒟 and its subsets 𝒟_10_, 𝒟_100_, 𝒟_1000_ and 𝒟_10000_).

We also constructed MSAs from three sequence datasets: UniRef90, UniRef100 and Metagenomics (described in Section 2.1). As represented in Fig. 1a, the LOR_*w*_-ΔΔ*G* Spearman correlation coefficients are almost identical irrespective of the sequence dataset used. The use of larger sequence datasets enriches the MSAs with homologous sequences. However, this neither increases nor decreases the correlation, which means that the additional sequences probably duplicate information already present in the MSAs constructed with the smaller UniRef90 sequence dataset. Note that this is different from what happens in protein structure prediction, where querying large metagenomics datasets for the input MSA construction seems to boost method performance [32, 33].

In addition to the factors mentioned above, the quality of an MSA also depends on the E-value threshold used to construct it. We tested E-value thresholds between 1 and 10^−22^, knowing that the default value in JackHMMER is 10^−3^. As shown in Fig. 1b, the LOR_*w*_-ΔΔ*G* correlation is relatively stable for E-values between 10^−2^ and 10^−8^. The highest correlation coefficient (0.467) is reached for an E-value threshold of 10^−7^. For E-value thresholds below 10^−13^, the correlation drops significantly, since the obtained MSAs become significantly smaller and composed of sequences that are very close to the query. Following the above analysis, in the rest of the article we will use MSAs constructed from UniRef90 with two JackHMMER iterations and an E-value threshold of 10^−7^.

We also evaluated the effect of MSA curation by removing sequences based on their gap ratio and sequence identity with the query sequence. We found almost no effect of the curation on the LOR_*w*_-ΔΔ*G* Spearman correlation coefficients, regardless of the sequence dataset used (see Supplementary Section 4). While MSA curation has been reported as an important step for better predictions [31], its effect is marginal on dataset 𝒟. This is probably due to the fact that 𝒟 only contains small protein domains, which makes MSA construction simpler. For instance, larger, multi-domain proteins usually have MSAs with sequences that only partially cover the query sequence, on which curation could have a stronger effect.

Finally, we leveraged the large-scale and systematic nature of the deep mutational scanning dataset 𝒟 to explore the impact of *N*_eff_, the effective number of sequences in an MSA, on the correlation between evolutionary scores and protein stability changes. While it is well known that the depth of an MSA increases its ability to make predictions [32, 31, 30] we are now able to more precisely quantify this relation and to further explore its application to protein stability. As shown in Fig. 2, MSAs with higher *N*_eff_ consistently lead to stronger LOR_*w*_-ΔΔ*G* correlations (and this holds true for other evolutionary models). This ensures that the MSA better represents the diversity of homologous proteins. Indeed, for the best E-value threshold (10^−7^), the average Spearman correlation coefficients between LOR_*w*_ and ΔΔ*G* are equal to 0.37 on 𝒟_10_, 0.44 on 𝒟_100_, 0.48 on 𝒟_1000_ and 0.50 on 𝒟_10000_. However, it is worth noting that there are substantial differences in performance between proteins, even between proteins with very similar *N*_eff_ values. The wide range of correlations (from 0.2 to 0.7) can thus not be fully attributed to *N*_eff_ alone. For instance, some proteins with very shallow MSAs, composed of only a few dozen sequences, are surprisingly predicted quite accurately by evolutionary models. This phenomenon is not limited to LOR_*w*_; proteins for which ΔΔ*G* is well predicted by one evolutionary model tend to be well predicted by the other models as well, and vice versa.

**Figure 2:**
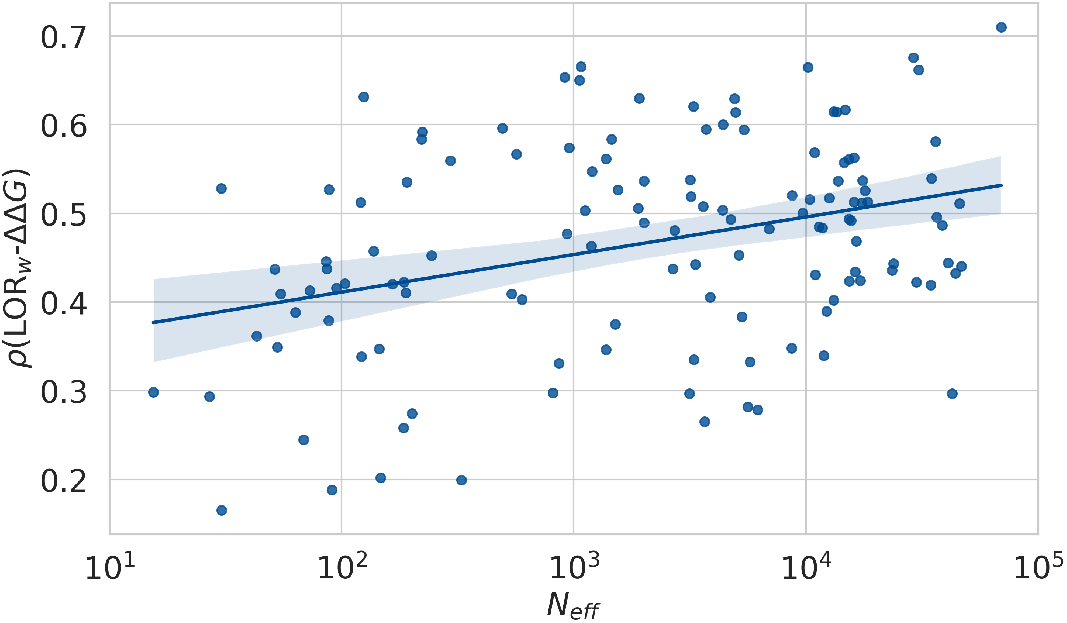
Per-protein Spearman correlation coefficients *ρ* as a function of the effective number of sequences *N*_eff_ (in log_10_ scale) in the MSAs (using UniRef90, two JackHMMER iterations and E-value threshold of 10^−7^).

### 3.2 Divergent predictions on highly similar proteins

To explore the strong variability between proteins in how well evolutionary models describe their stability, we focused on two proteins from 𝒟 with high sequence identity but notable differences in prediction performance. Both are engineered 57-residue *β*1 domains of the streptococcal G protein. The first, *Gβ*1_*pH*_, is a pH-sensitive mutant of *Finegoldia magna* whose sequence is identical to that of the PDB structure 2ZW1 [60] with the substitution V54S (called 2zw1 A 0-56 V54S in our dataset). The second, 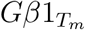, is a redesigned hyperthermophilic heptamutant from *Streptococcus sp. G148*; its sequence is identical to that of the PDB structure 1GB4 [61] with the substitution F53D (referenced as `1gb4_A_1-57_F53D`).

The two proteins share 70% sequence identity and very similar 3D structures, with a root mean square deviation (RMSD) of their C_*α*_ atoms of 0.4 Å (Fig. 3a). However, they show very different results in terms of LOR_*w*_-ΔΔ*G* Spearman correlation coefficients : *ρ* = 0.53 and 0.17 for *Gβ*1_*pH*_ and 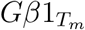, respectively (Tab. 1 and Fig. 3c-3d).

**Figure 3:**
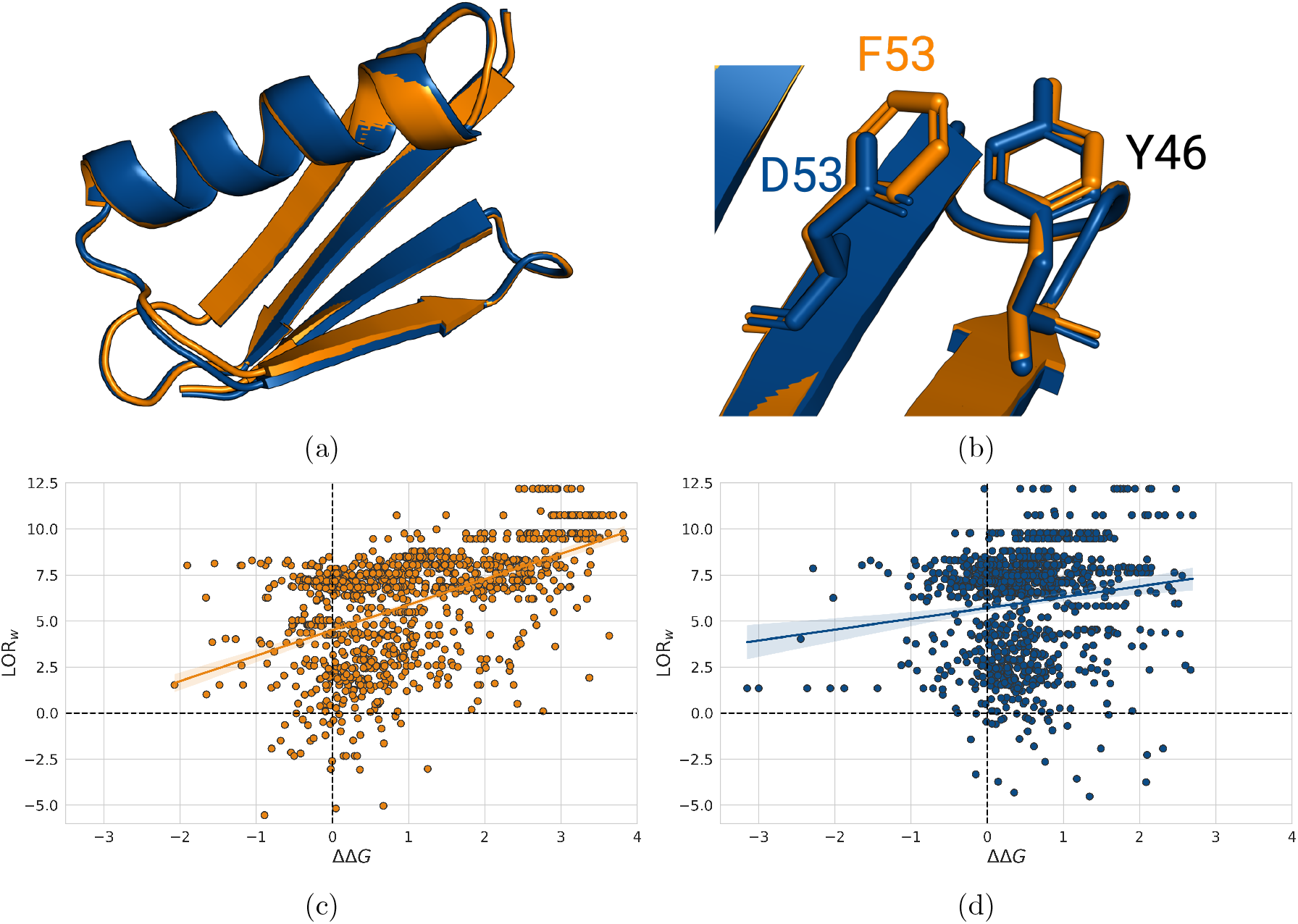
(a-b) Superimposed structures of proteins *Gβ*1_*pH*_ (in orange) and 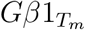 (in blue). (b) Interactions between residues 46 and 53. (c-d) Comparison between LOR_*w*_ scores and experimental ΔΔ*G* values for *Gβ*1_*pH*_ (c, in orange) and 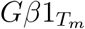 (d, blue).

**Table 1:** Spearman correlation coefficients *ρ* between experimental ΔΔ*G* values and scores obtained with LOR_*w*_, pycofitness (PCF) [53] and PoPMuSiC (PoP) [12] for the two G*β*1 proteins.

| Protein | Length (aa) | $N_{\text{eff}}$ | $\rho(LOR_w-\Delta\Delta G)$ | $\rho(PCF-\Delta\Delta G)$ | $\rho(PoP-\Delta\Delta G)$ |
| --- | --- | --- | --- | --- | --- |
| $G\beta 1_{pH}$ | 57 | 30.3 | 0.53 | 0.54 | 0.63 |
| $G\beta 1_{T_m}$ | 57 | 30.5 | 0.17 | 0.10 | 0.55 |

Since the two proteins have a close evolutionary relationship, they have almost identical MSAs and their evolutionary scores are expected to be similar. This is indeed the case: the Spearman correlation between their LOR_*w*_ values at positions sharing the same amino acid is equal to 0.99. Notably, however, their ΔΔ*G* scores at the same positions display a much lower Spearman correlation of 0.75. This suggests that epistatic effects, which are not captured by LOR_*w*_ scores, play an important role in the stability properties of these proteins.

An example of a particularly strong epistatic effect can be observed by looking at residue 46, which is occupied by a tyrosine in both *Gβ*1_*pH*_ and 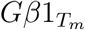. This position is highly conserved, with 98% of tyrosine in both MSAs; it thus has a strongly unfavorable LOR_*w*_ score for substitutions. However, while mutations at that position are all strongly destabilizing for *Gβ*1_*pH*_ (average ΔΔ*G* of 3.0 kcal/mol), their effect is only mildly destabilizing to neutral for 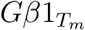. Analyzing this residue in its 3D structure, we found it to be located in a partially buried region and to interact with residue 53. It forms a stabilizing *π* − *π* (parallel-displaced) interaction with F53 in *Gβ*1_*pH*_, and an anion −*π* interaction with D53 in 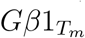, known to be barely stabilizing (Fig. 3b). Phenylalanine at position 53 is highly conserved in evolution, whereas aspartate at this position only occurs in the sequence of 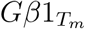. The F53D substitution from *Gβ*1_*pH*_ to 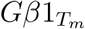 has thus a strong epistatic effect on position 46: in short, it changes Y46 mutations from very destabilizing to stabilizing or neutral, an effect that cannot be captured by independent-site LOR_*w*_ scores. As a consequence, LOR_*w*_ is less accurate for Y46 mutations in 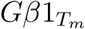 than in *Gβ*1_*pH*_.

Coevolution-based models that account for epistatic interactions should perform better and capture these effects. Unfortunately, this does not appear to be the case, as the mean pycofitness scores for all substitutions of Y46 are the same for both proteins. In addition, the Spearman correlations of ΔΔ*G* values with pycofitness scores are very similar to those with LOR_*w*_ scores for both proteins, as shown in Tab. 1, suggesting a difficulty in efficiently learning epistatic patterns. Similar results were observed for epistatic models other than pycofitness. We will return to the ability of such models to inform about protein stability in the next subsection.

Finally, we compared the results of evolutionary models with those of PoPMuSiC [12], a structure-based ΔΔ*G* prediction model that relies on statistically derived physical features. Although a small difference in performances is still observed between *Gβ*1_*pH*_ and 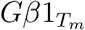, PoPMuSiC is able to better capture the effect of mutations in both proteins, with correlation values of 0.63 and 0.55, respectively.

These observations show that the ability of evolutionary information to explain stability differs from one protein to another. Although we were able to observe and analyze such differences on a per-protein basis, we were unable to identify biophysical protein characteristics or properties of their MSAs that could consistently explain this variability.

For example, although we interpreted that the low prediction accuracy of Y46 mutations in 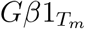 was due to the interacting residue at position 53 being occupied by an unusual amino acid, this can not be generalized to the entire sequence. Rather, it appears that a few key positions, such as 53, have complex epistatic effects on the mutational landscape and make the effects of mutations less predictable using only their evolutionary history [62]. Analyses performed on experimental deep mutational scanning data of domain *β*1 of protein G reveal that a minority of pairs (about 5%) exhibit significant epistatic effects on structural stability, but that a larger fraction (about 30%) exhibit weaker forms of epistasis [63] that are even more difficult to detect.

### 3.3 Assessing independent-site and epistatic evolutionary models for ΔΔ*G* prediction

We evaluated the ability of various evolution-based methods to detect the impact of mutations on protein stability. More precisely, we tested the independent-site models LOR, LOR_*w*_, GEMME-ind [31] and EVcouplings-ind [55], as well as the epistatic models pycofitness [53], ArDCA [56], GEMME-epi [31] and EVcouplings-epi [55]. For that purpose, we computed the average per-protein Spearman correlation coefficient between these evolutionary scores and experimental ΔΔ*G* values on dataset 𝒟 and on its subsets (split by MSA depth, see Section 2.2). The performance of the evolutionary models is shown in Fig. 4 and Tab. 2 (*p*-values for pairwise correlation comparisons and the method to derive them are provided in Supplementary Section 3).

**Figure 4:**
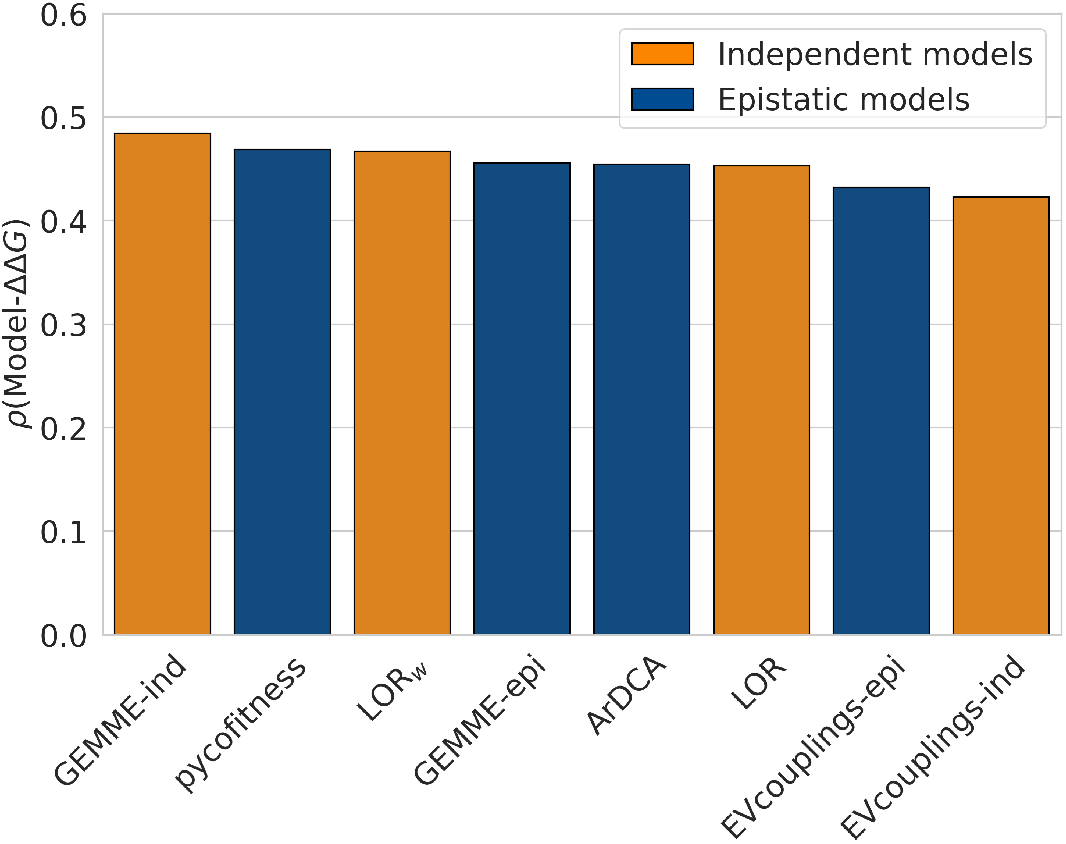
Average per-protein Spearman correlation coefficients *ρ* between experimental ΔΔ*G* values and evolutionary scores derived from the tested independent-site models (in orange) and epistatic models (in blue), on dataset 𝒟.

First, we observe that the performance is in general surprisingly good for evolutionary features, which are not always directly related to protein stability. Indeed, a low-frequency mutant residue in an MSA does not necessarily imply reduced protein stability, but can be caused by a different effect such as reduced function. Moreover, regardless of the method tested, the correlations are, on the average, better for proteins that have a large number of homologs, as expected. Correlation coefficients close to 0.5 are reached for all tested methods on dataset 𝒟_10000_. However, the performance remains surprisingly good, albeit lower, for shallow MSAs, with correlation coefficients in the 0.25 - 0.4 range on dataset 𝒟_10_. Most importantly, while epistatic models are more complete and have already established their relevance in multiple applications [64], simple independent-site methods appear to be equally predictive of stability changes (with GEMME-ind statistically significantly outperforming all epistatic models). Whether this trend is due to the relatively low impact of epistatic effects on stability or to the poor ability of these models to describe mutational effects remains to be understood.

In summary, evolutionary-based features are able to accurately estimate stability changes upon mutations. Independent-site models, which assume that residues have evolved independently of their context, tend to perform as well or even slightly better than more complex epistatic models when correlated to ΔΔ*G*, even though the latter take into account epistatic interactions that are known to play an important role in the evolutionary trajectory of proteins [25, 62, 65].

### 3.4 Improving performances of epistatic models

The surprising result of the previous section, that epistatic models, despite being more complex, do not perform better than simpler independent-site models, led us to further investigate the broader question of the role of covariation in predicting mutational effects. It is well known that in DCA methods, the inference of couplings often suffers from noise and sampling issues, which are reflected in the accuracy of the resulting coevolutionary models [66, 67]. Note that, while in contact predictions only highly coevolving pairs are considered, in studying the mutational landscape, in principle, all possible couplings are taken into account, thus amplifying the effect of potential noisy inference. We thus performed several checks to verify the aforementioned behaviors on our dataset. We assessed whether epistatic models are more sensitive to MSA depth than independent-site methods. As shown in Table 2, ArDCA appears to be much more sensitive to MSA depth than other models, with correlations as low as 0.25 for proteins with the shallow MSA (𝒟_10_) and as high as 0.53 for proteins with the largest MSA (𝒟_10000_). However, other epistatic and independent-site models show a similar sensitivity to MSA depth, and this feature alone cannot account for the fact that the more complex epistatic models do not outperform the simpler independent-site models. To further support these observations, we performed an MSA subsampling analysis, which confirmed ArDCA’s high sensitivity to MSA depth compared to other methods (see Supplementary Section 6.1).

**Table 2:**
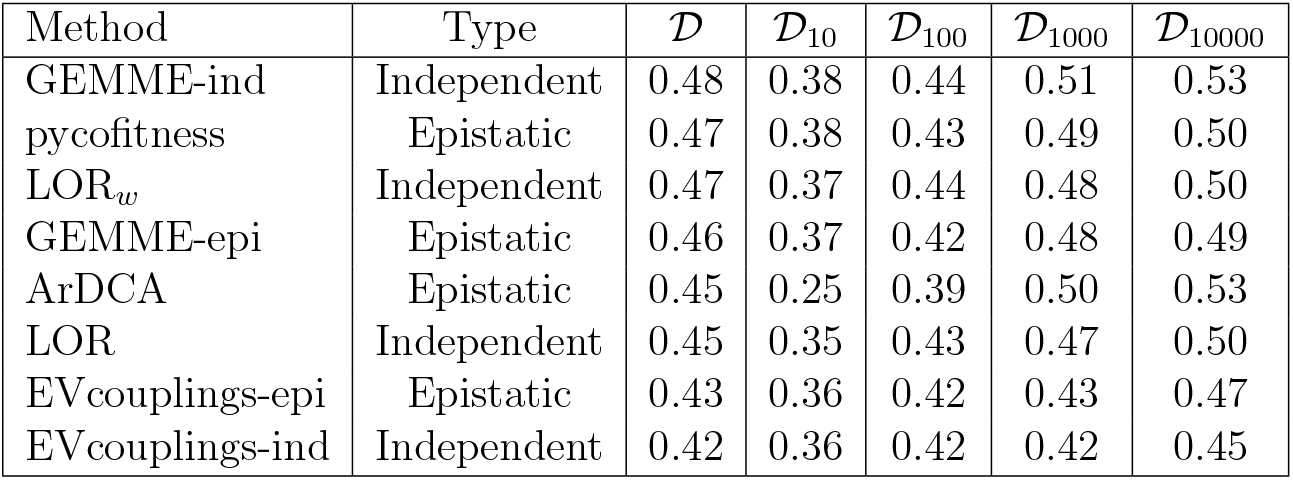
Average per-protein Spearman correlation coefficients *ρ* between experimental ΔΔ*G* values and evolutionary scores derived from independent-site and epistatic models, on dataset 𝒟 and its *N*_eff_-dependent subsets.

| Method | Type | $\mathcal{D}$ | $\mathcal{D}_{10}$ | $\mathcal{D}_{100}$ | $\mathcal{D}_{1000}$ | $\mathcal{D}_{10000}$ |
| --- | --- | --- | --- | --- | --- | --- |
| GEMME-ind | Independent | 0.48 | 0.38 | 0.44 | 0.51 | 0.53 |
| pycofitness | Epistatic | 0.47 | 0.38 | 0.43 | 0.49 | 0.50 |
| LOR<sub>w</sub> | Independent | 0.47 | 0.37 | 0.44 | 0.48 | 0.50 |
| GEMME-epi | Epistatic | 0.46 | 0.37 | 0.42 | 0.48 | 0.49 |
| ArDCA | Epistatic | 0.45 | 0.25 | 0.39 | 0.50 | 0.53 |
| LOR | Independent | 0.45 | 0.35 | 0.43 | 0.47 | 0.50 |
| EVcouplings-epi | Epistatic | 0.43 | 0.36 | 0.42 | 0.43 | 0.47 |
| EVcouplings-ind | Independent | 0.42 | 0.36 | 0.42 | 0.42 | 0.45 |

MSA curation is also important in epistatic model inference. For example, removing MSA columns with a high gap frequency can improve the performance of the inferred epistatic model, as it is expected to reduce the noise in the coupling parameters, while having no effect on independent-site models. To show this, we performed MSA trimming by removing columns with gap frequency higher than 0.2 and found an improvement of the epistatic model performances (results reported in Supplementary Section 6.2). This indicates that epistatic couplings are more sensitive than independent-site models to noise in MSAs.

When inferring an epistatic model, we also have to address the well-known issue of undersampling [68, 48, 67], as the number of parameters usually largely exceeds the number of sequences in the input MSA. Statistical regularization is commonly employed to prevent overfitting by adding penalty terms. These terms play an important role, as is well known in the DCA literature [67, 56, 30]. It has been shown that with different regularization strengths, the inference focuses on different features, shifting from local interactions to larger-scale functionally critical regions. [67]. In Supplementary Section 6.3, we show how the choice of the regularization parameters can enhance the ability of epistatic model to predict thermodynamic stability. Interestingly, we also found that different regularization strengths have opposite effects on predicting mutations in the core and on the surface of proteins. Weak regularization tends to improve core residue predictions, while strong regularization is optimal for surface residues. There is currently no clearly defined strategy for selecting the best regularization parameters to improve epistatic model performance. As all our analyses suggest that the epistatic models suffer from noisy couplings to the extent that we do not observe any significant advantage over independent-site methods, we further investigated this point by developing a new epistatic model that includes only a subset of couplings. A similar approach was employed in [66], where a supervised selection strategy was used to retain only couplings that enhance performance. In our model, we used an unsupervised approach that retains only coupling terms corresponding to highly coevolving residue pairs (see Supplementary Section 6.4). Specifically, we measured the coupling strength between two residue positions *i* and *j* by their Frobenius norm *F*_*ij*_, and only considered epistatic couplings between positions pairs whose Frobenius norm exceeds a given threshold. We found that this approach significantly improves model performance, as shown in Fig. 5, leading to better performances than independent-site models. More details can be found in Supplementary Section 6.4.

**Figure 5:**
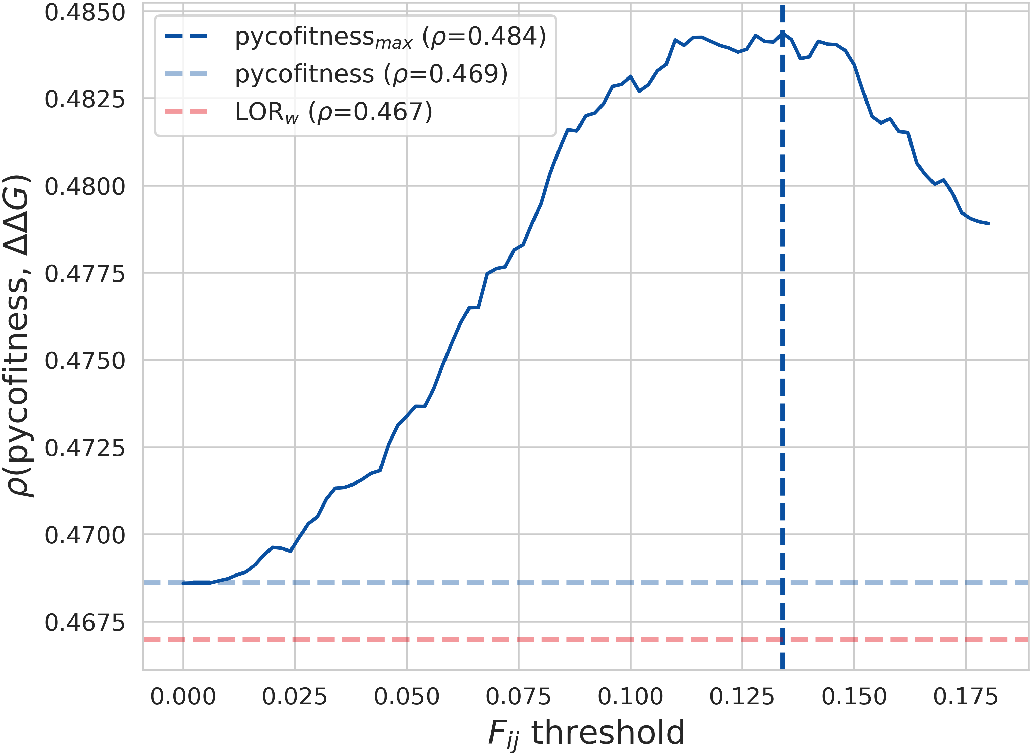
Average per-protein Spearman correlation coefficient *ρ* between the experimental ΔΔ*G* values and the Frobenius norm-modified pycofitness score as a function of the Frobenius norm threshold *t* on dataset 𝒟. Horizontal lines represent baseline comparisons: LOR_*w*_ score in red and default pycofitness score in blue. The vertical line represents the optimal Frobenius norm threshold *t*.

In summary, our results indicate that current full epistatic models do not improve performance in predicting the impact of mutations on thermodynamic stability compared to independent-site methods due to noisy couplings. However, note that, when predicting other functional properties, such as fitness, this seems to be less the case, as shown in previous analyses [69], which suggests that epistatic models may be better at capturing functional effects than stability.

### 3.5 Residue solvent accessibility and protein stability

It is well known that RSA plays an essential role in shaping protein stability, as substitutions in the core, where residues are more constrained by multiple interactions with surrounding residues, tend to be more destabilizing than those at the surface. As a result, many state-of-the-art stability change predictors (such as [12, 70, 18, 71]) directly or indirectly use RSA as a feature in their model. The large-scale and systematic nature of the deep mutational scanning dataset 𝒟 enables a more comprehensive exploration and quantification of this relationship. Furthermore, the analysis in this section offers an explanation of how combining RSA with evolutionary metrics enhances ΔΔ*G* predictions, highlighting that this effect is particularly pronounced for stability compared to other mutational properties like binding or activity.

We first emphasize the notable anti-correlation of −0.50 between RSA and ΔΔ*G*. As shown in Fig. 6a, mutations in low RSA regions display a wide distribution with high destabilizing median values, while mutations in high RSA regions display a narrow distribution with a median only slightly above zero kcal/mol.

**Figure 6:**
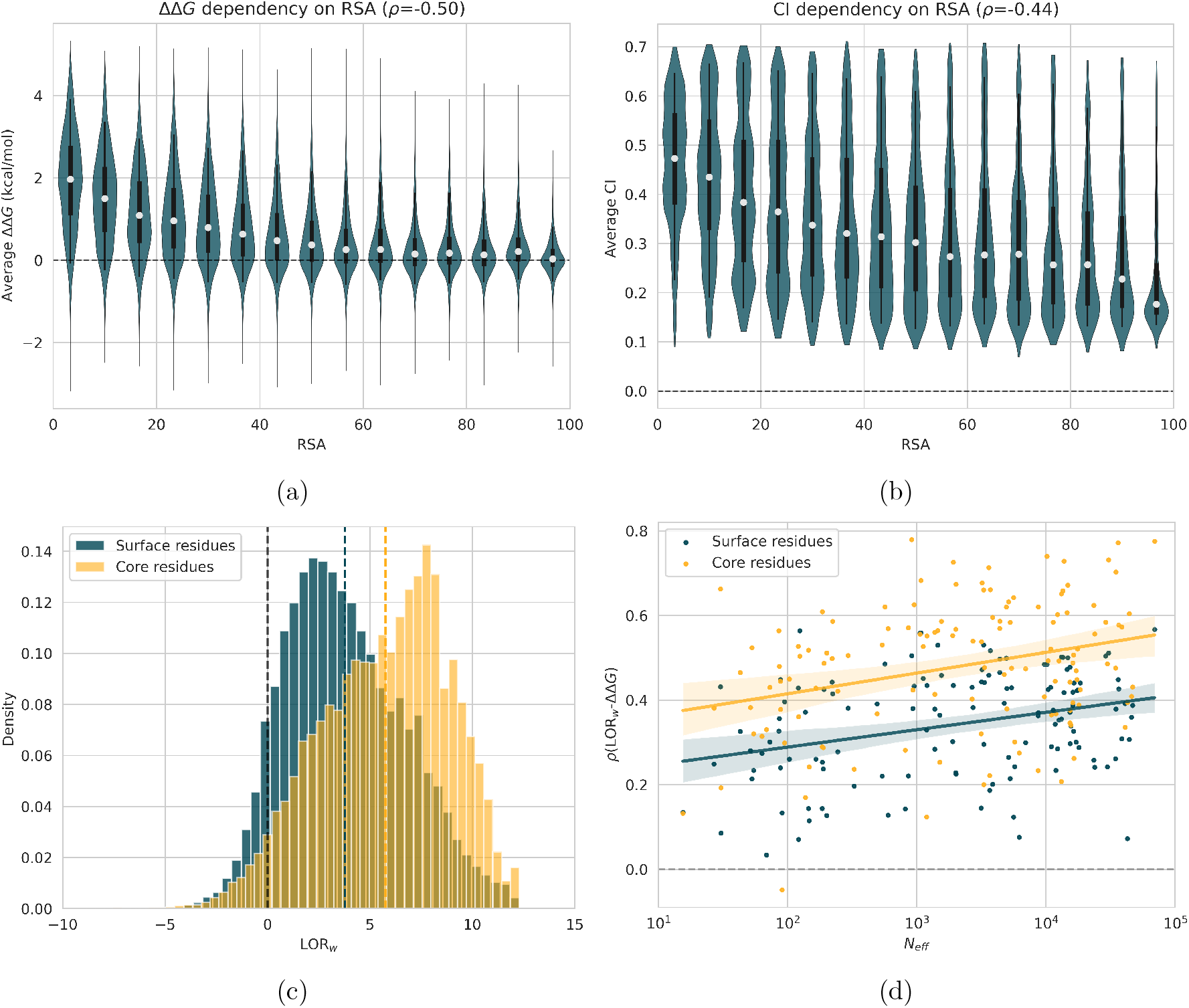
Distributions of (a) experimental ΔΔ*G* values in dataset 𝒟 and (b) corresponding CI values as a function of the RSA of the wild-type residues. White dots represent the median, boxes (thick black lines) represent the inter-quartile range from quartile 1 to quartile 3, and whiskers (thin back lines) represent the inter-percentile range from percentile 5 to percentile 95. (c) Distribution of LOR_*w*_ for mutations located in the core (in orange) and at the surface (in blue). (d) Per-protein Spearman correlation coefficients *ρ* between LOR_*w*_ and experimental ΔΔ*G* values for mutations located in the core (in orange) and at the surface (in blue) as a function of *N*_eff_ (in log_10_ scale) in the corresponding MSA.

The RSA dependence is particularly strong for the effect of variants on stability. This can be seen by examining the RSA dependence of single-site mutation scores from ProteinGym [72], a large-scale standard dataset on the effect of protein variants on fitness, which collects many multi-label deep mutational scanning experiments. RSA shows an average per-protein Spearman correlation of −0.50 on mutagenesis data from stability-based assays (essentially composed of values from [16] similar to our dataset 𝒟). In contrast, data from other type of assays exhibit still remarkable but much weaker average correlations equal to −0.27, −0.28, −0.32 and −0.33 for binding-, organismal fitness-, expression- and activity-based assays, respectively.

We note that the observed anti-correlation between RSA and ΔΔ*G* may be overestimated on the dataset 𝒟 because it contains only small monomeric protein domains. In general, larger proteins have a larger fraction of their residues with near-zero RSA values. To test this, we analyzed the 707 mutations from the ℒ dataset, inserted in proteins ranging in length from 108 to 452 residues. We found that the average per-protein Spearman correlation between RSA and ΔΔ*G* is only slightly smaller, with a value of −0.42 (see Supplementary Section 2 for details).

Values of RSA are also strongly related to evolution, as residues at the surface tend to evolve much faster than those in the core [73, 74, 75]. This is confirmed by the anti-correlation of −0.44 between RSA and the conservation index CI (Fig. 6b). Unconserved average values for high RSA regions are explained by the fact that such residues are mostly located in variable loop regions or at the C- and N-terminus of the protein, where the evolutionary pressure is low compared with the higher evolutionary pressure that is exerted on core residues. From the perspective of the mutational evolutionary score LOR_*w*_, we found its average value to be notably higher for core residues (RSA *<* 20%) than for surface residues (RSA ≤ 20%) with averages of 5.75 and 3.78, respectively, as shown in Fig. 6c. Moreover, LOR_*w*_ displays a stronger predictive power on ΔΔ*G* for core residues than for surface residues (*ρ* = 0.48 and *ρ* = 0.34 respectively, see Fig. 6d). However, other factors may influence this result. First, surface residues are submitted to strong evolutionary pressures from other protein properties such as the optimization of the binding to other proteins, ligands or nucleic acids. Second, since surface residues display smaller stability variations upon mutations, these effects are more subtle and thus more difficult to predict.

### 3.6 Combining RSA and evolution for ΔΔ*G* prediction

As shown in the previous subsection, RSA is strongly related to both protein stability and conservation. The higher evolutionary pressure for stability in the core of proteins makes RSA a promising indicator to modulate signals extracted from MSAs. We therefore used the RSA as defined in Eq. 11 in order to improve the predictive power of the evolutionary scores for ΔΔ*G* prediction. In addition, the RSA factor helps to mitigate predictions in MSA regions with a high gap ratio. Indeed, protein loop regions and the C- and N-termini tend to be poorly conserved, leading to variable MSAs with many gaps. While mutations in these flexible and exposed regions are often associated with neutral ΔΔ*G* values, they tend to be incorrectly classified as destabilizing by evolutionary models due to the lack of specific sequence information in these regions. Inclusion of the RSA factor helps to lower the predicted scores, which then better describe the tendency of these mutations to be neutral.

The performance of all evolutionary methods combined with RSA to predict protein stability changes on the dataset 𝒟 is presented in Fig. 7; the correlation values are provided in Supplementary Section 7, and the *p*-values for pairwise correlation comparisons are given in Supplementary Section 3. Additionally, we compared them to seven state-of-the-art ΔΔ*G* prediction models (PoPMuSiC [12], KORPM [76], MAESTRO [70, 77], RaSP [78], PremPS [18], DDMut [71], and DDGun3D [14]). The first four predictors are purely structure-based; they rely solely on biophysical and statistical features and identify the parameters of their models based on experimental ΔΔ*G* datasets. In contrast, the last three methods include evolutionary features in addition to structure-based features. PremPS extracts them from MSAs, whereas DDGun3D and DDMut use general evolutionary metrics such as mutation matrices and sequence-based statistical potentials. Notably, DDGun3D is almost unsupervised as it only uses experimental ΔΔ*G* datasets to set the weight coefficient between structural and evolutionary features. Also, only RaSP and DDMut use complex deep learning techniques. It is interesting to observe the emergence of transformer-based protein large language models (pLLMs) such as PROSTATA [79] and Stability Oracle [80], which show promising performances. However, their learning sets partially rely on the deep mutational scanning dataset [16] from which we constructed our test dataset 𝒟, and therefore these methods cannot be used for a fair comparison in our analysis.

**Figure 7:**
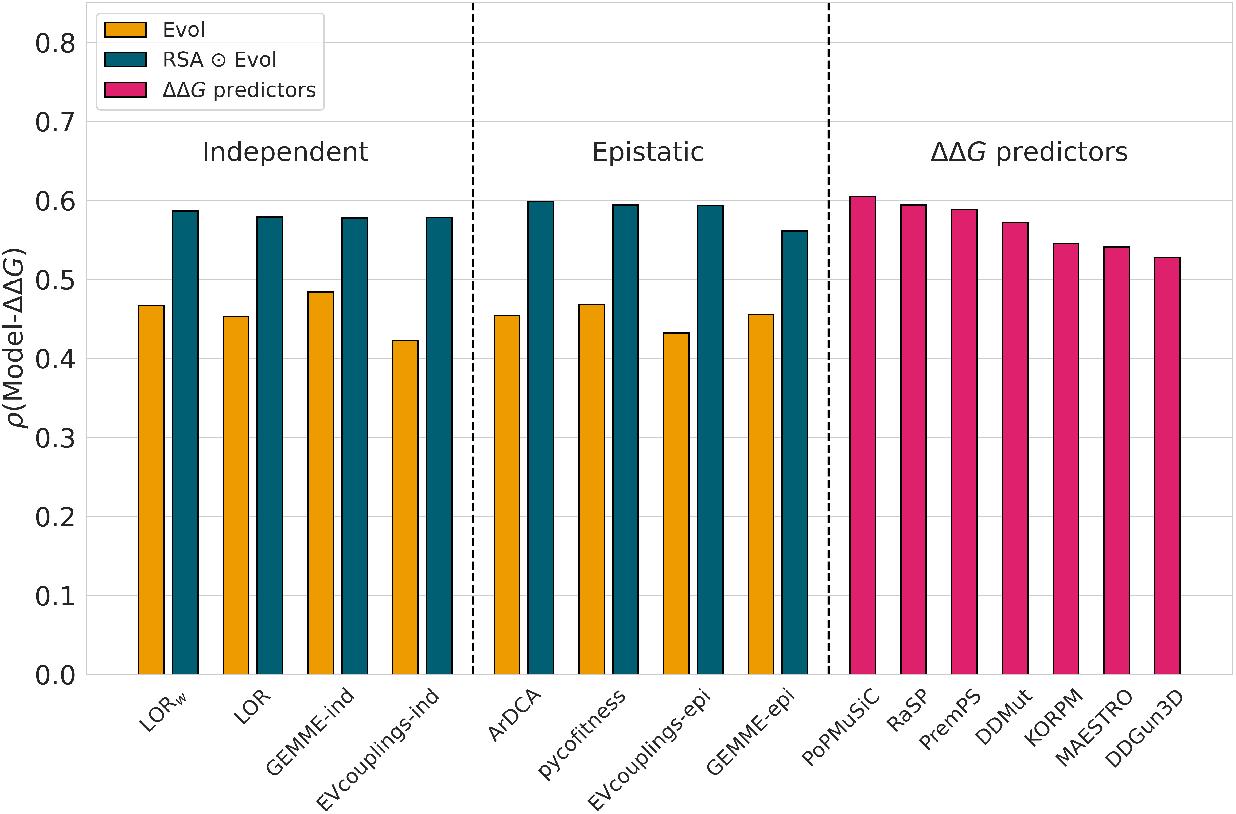
Average per-protein Spearman correlation coefficients *ρ* between experimental ΔΔ*G* values and evolutionary scores (in yellow), evolutionary scores combined with RSA (in dark blue) and predicted ΔΔ*G* values (in pink), on the variant dataset 𝒟.

Modulation by RSA significantly improves the performance of all evolutionary models, increasing the correlations from *ρ* in the range [0.42, 0.48] up to [0.56, 0.60]. Surprisingly, these scores are comparable or in some cases even better than those of the supervised ΔΔ*G* prediction models. For example, the best evolutionary score of 0.60 achieved by RSA ⊙ ArDCA outperforms nearly all ΔΔ*G* predictors and is statistically comparable to the three top-performing predictors (PoP-MuSiC, RaSP, and PremPS with no statistically significant difference in *p*-values). As shown in Supplementary Section 2, this high correlation is also confirmed for mutations in larger proteins from the ℒ dataset, with an average per-protein Spearman correlation of 0.58 for RSA ⊙ LOR_*w*_.

Overall, independent-site and epistatic models achieve comparable performance when modulated by RSA. However, we note that in this case, three epistatic methods (ArDCA, pycofitness, and EVcouplings-epi) outperform all independent-site methods, with relatively small but statistically significant differences (especially on proteins with large *N*_eff_). In terms of MSA depth, the RSA-modulated models, as expected, behave similarly to the evolutionary models alone, with performance improving as MSA depth increases (see Supplementary Section 7). For shallow MSAs, their performance is only slightly better than RSA alone, indicating the weak quality of the extracted (co)evolutionary signals. In contrast, for deeper MSAs, the performance is significantly higher.

In conclusion, in the era of deep learning and large language models, it is impressive to see that simple unsupervised models including only amino acid frequencies and their localization in the structure achieve state-of-the-art performance.

## 4 Conclusion

Evolution and protein stability are two correlated quantities. Indeed, protein stability is known to be one of the major factors driving evolution, and on the other hand, evolution puts constraints on protein stability. In this paper, we took advantage of the huge mutational stability dataset published in [16] to carefully analyze how to leverage evolutionary information to predict the effect of mutations on protein stability. The main findings of the paper are summarized below:

- The quality of the used MSA plays a fundamental role in the ability of evolution-based methods to extract information about protein stability. In particular, proteins with deeper MSAs tend to be largely better predicted than proteins with shallow MSAs. However, having more sequence data does not necessarily lead to better scores, as already highlighted in [37].
- Among the various evolutionary models tested to predict the effect of mutations on protein stability, we found that independent-site models perform as well as or slightly better than coevolutionary models. Thus, although these models achieve very good performance in predicting 3D contacts between residues, they do not seem to have an advantage over independent-site models in predicting changes in protein stability.
- Due to the complexity of model inference, coevolutionary models often have noisy couplings. Various strategies, including MSA curation, parameter selection, and the choice of a subset of couplings, can be employed to improve the performance of epistatic models.
- RSA is a key feature strongly anti-correlated with changes in protein stability caused by mutations. Even more impressively, the simple unsupervised combination of RSA with the evolutionary features we analyzed leads to performance similar to that of the state-of-the-art supervised ΔΔ*G* predictors.

In summary, we have demonstrated that evolutionary data can be effectively used in an unsupervised way to predict the impact of mutations on protein stability. Since the performance obtained by modulating evolutionary signals with RSA is similar to that of complex, supervised ΔΔ*G* predictors, an immediate perspective is to combine these models with the aim of further improving ΔΔ*G* prediction.

## Acknowledgment

We acknowledge financial support from the Belgian Fund for Scientific Research (F.R.S.-FNRS) through a PDR project. M.T. benefits from a FNRS-FRIA PhD grant. P.H. benefits from a Win4Doc grant from SPW Recherche of the Walloon Region.

