## Supplementary Material for "Exploring evolution to uncover insights into protein mutational stability"

November 7, 2024

#### Contents

**1 Stability changes upon mutations in the DMS dataset $\mathcal{D}$ 2**

**2 Performances of evolutionary models to predict $\Delta\Delta G$ on S4038 3**

**3 $P$-value estimation of pairwise correlation comparisons 4**

**4 Impact of MSA construction and curation on evolutionary scores 6**

**5 Performance variation between proteins 7**

**6 Independent-site versus epistatic models: an in-depth comparison 8**

**7 Performances of evolutionary models to predict $\Delta\Delta G$ on $\mathcal{D}$ subsets 12**

### 1 Stability changes upon mutations in the DMS dataset $\mathcal{D}$

We represent in Fig. S1 the distribution of experimental $\Delta\Delta G$ values from the variant dataset $\mathcal{D}$, which were obtained through deep mutational scanning (DMS) experiments on 129 small protein domains. This dataset is a subset of single-site mutations from the massive experimental dataset [1], from which we have removed data on proteins that are too close in sequence identity or whose multiple sequence alignments (MSAs) were too small (see section 2.2 of the main text for details). Since protein stability exerts an evolutionary pressure on protein sequences and native structures generally correspond to an absolute free energy minimum, most of the mutations are destabilizing, as expected. The average $\Delta\Delta G$ value is equal to 0.82 kcal/mol.

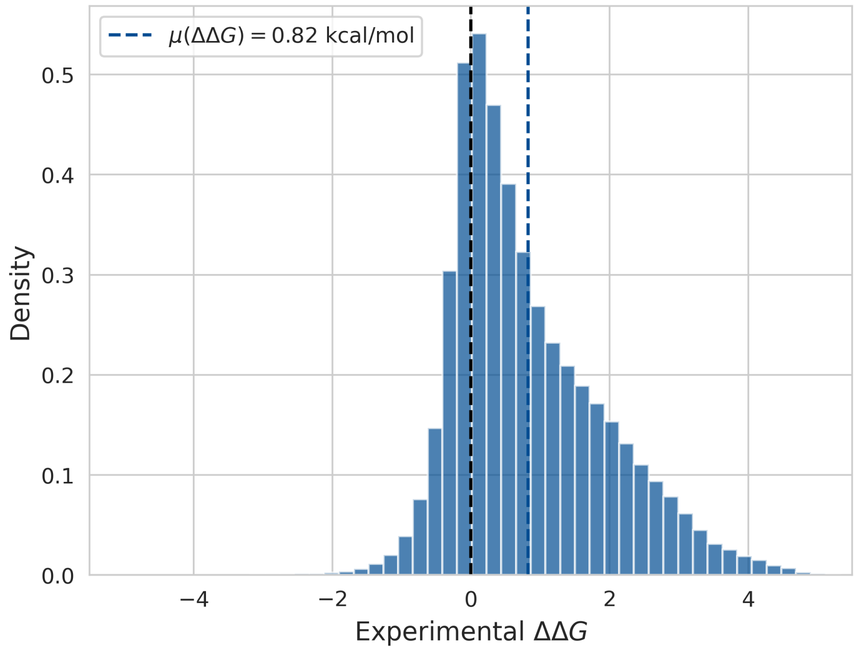

Figure S1: Distribution of experimental $\Delta\Delta G$ values (in kcal/mol) from the variant dataset $\mathcal{D}$ containing $135,056$ mutations.

#### 2 Performances of evolutionary models to predict $\Delta\Delta G$ on S4038

Since our main dataset $\mathcal{D}$ only consists of small protein domains (about 60 residues), we wanted to evaluate how evolutionary information extracted from MSAs and its combination with RSA performs on larger proteins. For that purpose, we used the recently published literature-based dataset S4038 [2], even though it does not represent a systematic set of mutations. Note that its $\Delta\Delta G$ values were obtained with methods such as differential scanning calorimetry which are more accurate than the cDNA display proteolysis approach used to construct $\mathcal{D}$. However, its literature-based nature and the heterogeneity of the experimental conditions are likely to increase both systematic and random errors and create unbalances in the dataset [3].

From the S4038 dataset, we only included proteins with at least 60 mutations to reduce sampling biases in the selection of mutations and to be able to derive reliable per-protein Spearman correlations. Additionally, we selected only proteins with 100 residues at least, as our goal was to extend our results to longer proteins. The final dataset $\mathcal{L}$ consists of 707 mutations in 7 different proteins with lengths ranging from 108 to 452 residues.

We show in Fig. S2 and Tab. S1 the per-protein Spearman correlations between $\Delta\Delta G$ and the independent-site evolutionary feature $\mathrm{LOR}_w$ and its structure-informed counterpart $\mathrm{RSA} \odot \mathrm{LOR}_w$. We observe that, despite the larger protein lengths, the results on dataset $\mathcal{L}$ are in line with our observations on $\mathcal{D}$, with possibly a slightly smaller impact of the RSA on the performance. Indeed, the average correlation for $\mathrm{LOR}_w$ is even higher for dataset $\mathcal{L}$ than for $\mathcal{D}$, whereas for $\mathrm{RSA} \odot \mathrm{LOR}_w$, the scores are very similar. This is explained by the slightly higher anti-correlation between RSA and $\Delta\Delta G$ values in $\mathcal{D}$ than in $\mathcal{L}$. We cannot strictly exclude that very large multi-domain proteins may behave somewhat differently, given the lack of currently available data. However, Fig. S2 provides further evidence that our findings can be safely extended to longer proteins, as short and long proteins show similar Spearman correlation trends as a function of $N_{\text{eff}}$. Indeed, we observe non-significant $p$-values for the Mann-Whitney $U$ test between the correlations of $\mathcal{L}$ and $\mathcal{D}_{1000} \cup \mathcal{D}_{10000}$ (0.35 and 0.26 for $\mathrm{LOR}_w$ and $\mathrm{RSA} \odot \mathrm{LOR}_w$, respectively).

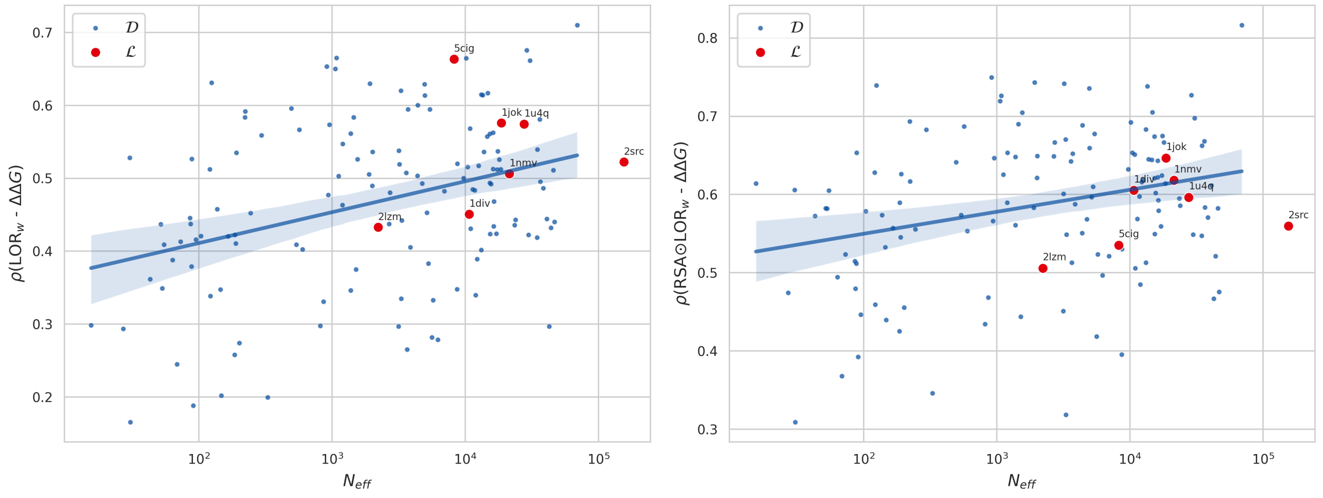

(a) (b)

Figure S2: Per-protein Spearman correlation coefficient $\rho$ as a function of $N_{\text{eff}}$ (in $\log_{10}$ scale) for the datasets $\mathcal{D}$ (blue dots with linear regression line) and $\mathcal{L}$ (red dots with proteins' PDB code) between: (a) $\Delta\Delta G$ and $\mathrm{LOR}_w$; (b) $\Delta\Delta G$ and $\mathrm{RSA} \odot \mathrm{LOR}_w$.

Table S1: Performance of $\text{LOR}_w$ and $\text{RSA} \odot \text{LOR}_w$ in predicting the $\Delta\Delta G$ values of mutations in the proteins contained in dataset $\mathcal{L}$, compared to their performance on dataset $\mathcal{D}$. Column 1 contains the PDB code of the mutated protein. Columns 2-4 show the Spearman correlations between the features RSA, $\text{LOR}_w$ and $\text{RSA} \odot \text{LOR}_w$, and the experimental $\Delta\Delta G$ values. $N_{\text{muts}}$ represents the number of mutations in the given proteins, and $N_{\text{tot}}$ and $N_{\text{eff}}$ denote the total and effective number of sequences in the MSA, respectively. "seq-len" is the length of the protein and "spread" is the maximum sequence separation between mutations in the protein.

| PDB | RSA | $\text{LOR}_w$ | $\text{RSA} \odot \text{LOR}_w$ | $N_{\text{muts}}$ | $N_{\text{tot}}$ | $N_{\text{eff}}$ | seq-len | spread |
|---|---|---|---|---|---|---|---|---|
| 2lzm | -0.50 | 0.43 | 0.51 | 214 | 3,882 | 2,214 | 164 | 161 |
| 5cig | 0.14 | 0.66 | 0.54 | 60 | 16,447 | 8,225 | 108 | 102 |
| 1div | -0.64 | 0.45 | 0.61 | 81 | 19,944 | 10,710 | 149 | 145 |
| 1jok | -0.71 | 0.58 | 0.65 | 150 | 32,267 | 18,649 | 149 | 146 |
| 1nmv | -0.39 | 0.51 | 0.62 | 68 | 108,353 | 21,429 | 163 | 68 |
| 1u4q | -0.43 | 0.57 | 0.60 | 74 | 220,769 | 27,672 | 322 | 204 |
| 2src | -0.38 | 0.52 | 0.56 | 60 | 1,243,130 | 154,997 | 452 | 109 |
| Average on $\mathcal{L}$ | -0.42 | 0.53 | 0.58 | 101 | - | - | - | - |
| Average on $\mathcal{D}$ | -0.50 | 0.47 | 0.59 | 1047 | - | - | - | - |

#### 3 $P$-value estimation of pairwise correlation comparisons

In this work, many performance scores were reported as per-protein average Spearman correlation coefficients in $\mathcal{D}$. Since the resulting values are often very close, we used a bootstrap method to determine whether the score from method $E_1$ is significantly higher than that from method $E_2$. However, we emphasize that the reported $p$-values should be interpreted with caution. In our case, even a very small difference in correlation can result in a statistically significant $p$-value due to the large size of dataset $\mathcal{D}$. Therefore, a statistically significant $p$-value indicates that methods $E_1$ and $E_2$ perform differently on $\mathcal{D}$, but does not reflect the magnitude of the difference nor does it guarantee that one method outperforms the other on a different dataset.

First, we constructed $\mathcal{D}_{ran}$ by randomly selecting, with replacement, 129 proteins from the 129 proteins in $\mathcal{D}$. Next, we computed the scores of $E_1$ and $E_2$ on this randomly sampled dataset $\mathcal{D}_{ran}$. The $p$-value was estimated as the fraction of sampled datasets $\mathcal{D}_{ran}$ for which the score of $E_1$ is higher than that of $E_2$, based on $100,000$ repetitions. It is important to note that, because correlation values vary significantly from protein to protein but tend to be close between different methods applied to the same protein (as discussed in Supplementary Section 5), each comparison has to be done on the exact same sampled dataset $\mathcal{D}_{ran}$.

We applied this bootstrap procedure to compare the performance of $\text{LOR}_w$ in predicting $\Delta\Delta G$ using MSAs derived with different JackHMMER parameters and sequence datasets (described in Section 3.1 of the main text) and found that:

- The use of different sequence datasets (UniRef90, UniRef100, and Metagenomics) does not lead to any statistically significant differences, except for a $p$-value of 0.03 indicating that UniRef100 performs slightly better than UniRef90 when the number of JackHMMER iterations is $n = 3$.
- When using UniRef90, varying the number of JackHMMER iterations ($n = 1, \ldots, 7$) shows that the optimal number is 2; this value is statistically significantly better than all other tested values.

We also applied this bootstrap procedure to compare the performance of different evolutionary metrics and $\Delta\Delta G$ predictors (described in Sections 3.3 and 3.5 of the main text):

- All evolutionary scores combined with RSA ($\text{RSA} \odot E$) outperform the individual score $E$, with a $p$-value of zero.

- When comparing the performance of the six studied evolutionary metrics, we observe that pycofitness is not significantly better than $LOR_w$, and that GEMME-epi is not significantly better than ArDCA and LOR. All other pairwise comparisons show statistically significant differences.
- When comparing the performance of the six evolutionary metrics combined with RSA, we found that pycofitness is not significantly better than EVcouplings-epi, and that LOR is not significantly better than EVcouplings-ind and GEMME-ind. All other pairwise comparisons show statistically significant differences.
- When comparing the performance of $\Delta\Delta G$ predictors, we observe that RaSP is not significantly better than PremPS, and that KORPM is not significantly better than MAESTRO. All other pairwise comparisons show statistically significant differences.
- We observe that the best evolutionary score combined with RSA, which is ArDCA, performs neither significantly better nor significantly worse than the best-performing $\Delta\Delta G$ predictors, i.e. PoPMuSiC, RaSP, and PremPS.

All pairwise $p$-values discussed above are available in our GitHub repository `https://github.com/3BioCompBio/EvoStability`.

### 4 Impact of MSA construction and curation on evolutionary scores

For all proteins of our variant dataset $\mathcal{D}$, MSAs were built using different sequence datasets (UniRef90, UniRef100 and Metagenomics defined in Section 2.1 of the main paper) and with different numbers of JackHMMER iterations. The Spearman correlation coefficients between the independent-site evolutionary features derived from these MSAs (i.e. LOR and $\text{LOR}_w$) and the experimental $\Delta\Delta G$ values are presented in Tab. S2. Consistently, $\text{LOR}_w$ performs slightly better than LOR. The optimal number of JackHMMER iterations was found to be 2, and no notable improvement is observed when increasing the sequence dataset from UniRef90 to the much larger datasets UniRef100 and Metagenomics.

Table S2: Average per-protein Spearman correlation coefficients $\rho$ between the experimental $\Delta\Delta G$ values and evolutionary features (LOR and $\text{LOR}_w$) on the variant dataset $\mathcal{D}$, as a function of MSA construction parameters, i.e. the number of JackHMMER iterations and the sequence dataset. The JackHMMER E-value threshold is set equal to $10^{-7}$.

| Sequence dataset | JackHMMER Iterations | LOR | $\text{LOR}_w$ |
|---|---|---|---|
| UniRef90 | 1 | 0.44 | 0.45 |
| UniRef90 | 2 | **0.45** | **0.46** |
| UniRef90 | 3 | 0.45 | 0.45 |
| UniRef90 | 4 | 0.44 | 0.45 |
| UniRef90 | 5 | 0.43 | 0.44 |
| UniRef90 | 6 | 0.43 | 0.44 |
| UniRef90 | 7 | 0.43 | 0.44 |
| UniRef100 | 1 | 0.43 | 0.45 |
| UniRef100 | 2 | **0.45** | **0.46** |
| UniRef100 | 3 | 0.44 | 0.46 |
| Metagenomics | 1 | 0.44 | 0.45 |
| Metagenomics | 2 | **0.45** | **0.46** |
| Metagenomics | 3 | 0.45 | 0.45 |

We also tested the effect of MSA curation on the performances of evolutionary score $\text{LOR}_w$ to predict $\Delta\Delta G$. Namely, we tested whether removing some sequences from the MSA can improve the correlation, as done in [4]. Fixing the number of JackHMMER iterations to two, we curated the MSAs obtained from the three sequence datasets (UniRef90, UniRef100 and Metagenomics) on the basis of sequence identity and gap ratio. For each sequence in the MSA that is not the wild-type sequence, the sequence identity and gap ratio were calculated as follows:

- The sequence identity (SI) is the number of identical aligned amino acids between the considered sequence and wild-type sequence, divided by the length of the wild-type sequence. Sequences most similar to the wild-type have a sequence identity close to 1, while the more distant ones have a score close to 0.
- The gap ratio (GR) is the fraction between the number of gaps (indels) in the considered sequence aligned to the wild-type sequence and the length of the wild-type sequence. Sequences containing the greatest number of gaps will have a high gap ratio, while a sequence containing no gaps will have a gap ratio of 0.

The Spearman correlation coefficients between the experimental $\Delta\Delta G$ values and the $\text{LOR}_w$ scores derived from these curated MSAs are presented in Tab. S3. We observe no notable improvement with different MSA curation methods, with only a marginal improvement when excluding sequences that are too far from the target sequence (sequence identity lower than 0.2).

Table S3: Per-protein Spearman correlation coefficients $\rho$ between the evolutionary score $\text{LOR}_w$ and the experimental $\Delta\Delta G$ values on the dataset $\mathcal{D}$. Sequences in the MSAs were filtered based on sequence identity (SI) and gap ratio (GR) criteria.

| Filter on MSAs' sequences | UniRef90 | UniRef100 | Metagenomics |
|---|---|---|---|
| Full MSA | 0.46 | 0.46 | 0.46 |
| Only sequences with $0.2 < \text{SI}$ | 0.46 | 0.46 | 0.46 |
| Only sequences with $\text{SI} < 0.8$ | 0.46 | 0.46 | 0.46 |
| Only sequences with $0.2 < \text{SI} < 0.8$ | 0.46 | 0.46 | 0.46 |
| Only sequences with $\text{GR} < 0.5$ | 0.46 | 0.46 | 0.46 |
| Only sequences with $\text{GR} < 0.2$ | 0.46 | 0.46 | 0.46 |
| Only sequences with $\text{GR} < 0.1$ | 0.45 | 0.45 | 0.45 |

### 5 Performance variation between proteins

The predictive power of evolutionary methods widely varies from one protein to another, with Spearman correlation coefficients ranging from less than 0.2 to more than 0.7, even for similar MSA depths. The MSA depth alone is therefore unable to explain this strong variability. To illustrate this point, we ranked the 129 proteins from dataset $\mathcal{D}$ according to how well each of the evolutionary methods predicts $\Delta\Delta G$ (using the Spearman correlation coefficient $\rho$). As shown in Fig. S3a, the rankings arising from different evolutionary methods are surprisingly similar (with the smallest but still very strong correlation of $\rho = 0.71$ between the GEMME-epi and EVcouplings-ind rankings). In contrast, the protein rankings by the different methods are much less correlated with the MSA $N_{\text{eff}}$ score; it ranges from 0.20 to 0.52 for EVcouplings-ind and ArDCA, respectively (Fig. S3b).

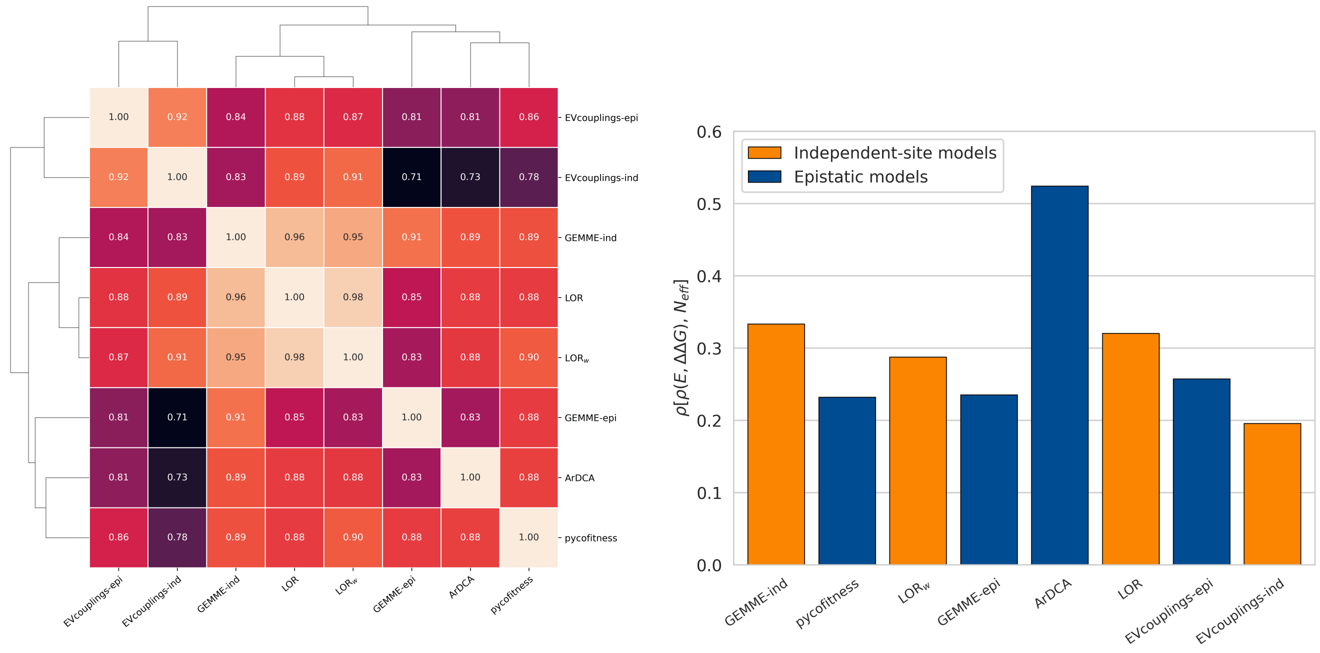

(a) (b)

Figure S3: For each evolutionary model, proteins from the dataset $\mathcal{D}$ are ranked according to how well the model scores are correlated with the experimental $\Delta\Delta G$ values (using the Spearman correlation coefficient). (a) Spearman cross-correlations between rankings of proteins according to different predictors. (b) Spearman correlations between $N_{\text{eff}}$ and the protein rankings.

### 6 Independent-site versus epistatic models: an in-depth comparison

We showed that, although epistatic co-evolutionary methods are theoretically more comprehensive, they fail to significantly outperform simpler independent-site approaches. This result is counterintuitive given the proven relevance of epistatic models in various fields [5, 6, 7, 8, 9, 10]. To investigate this discrepancy, we conducted analyses to identify the factors limiting the performance of epistatic models and explored potential strategies to address them.

#### 6.1 Robustness against subsampling

To investigate how different methods are affected by MSA depth (and in particular the apparent high sensitivity of ArDCA), we estimated the effect of random subsampling of MSAs on the performance of two independent-site models, $\mathrm{LOR}_w$ and EVcouplings-ind [6, 11], and two epistatic models, ArDCA [12] and EVcouplings-epi [6, 11]. From our dataset $\mathcal{D}$, we selected the 16 proteins displaying the deepest MSAs with $N_{\mathrm{eff}} > 20000$. We then subsampled the raw MSA by keeping only a fraction of all the initial sequences (randomly selected). We performed the analysis with a $N_{\mathrm{tot}}$ ratio ranging from $\frac{1}{2^1}$ to $\frac{1}{2^{12}}$, where this ratio is defined as the ratio between subsampled MSA depth and initial MSA depth. For more stable results, three independent random subsamplings were performed. Scores of the four evolution-based models were derived on the subsampled MSAs. As expected, the performance of all scores decreases with subsampling, but ArDCA displays a much larger decrease in performance and is thus more sensitive to MSA depth than the other three models (Fig. S4).

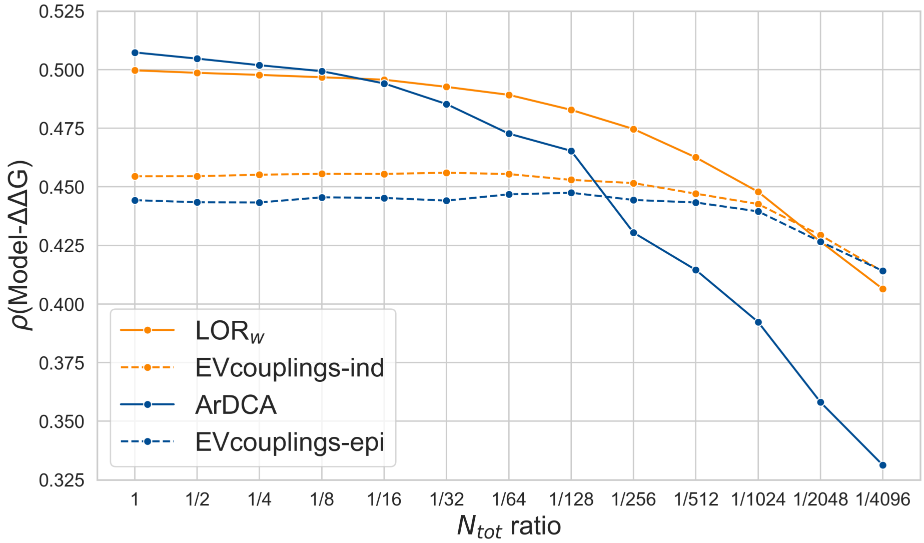

Figure S4: Average per-protein Spearman correlation coefficient $\rho$ between experimental $\Delta\Delta G$ values and $\mathrm{LOR}_w$, EVcouplings-ind, ArDCA and EVcouplings-epi scores as a function of the subsampling ratio $N_{\mathrm{tot}}$ (in negative logarithmic scale on the 16 proteins from $\mathcal{D}$ with the deepest initial MSAs).

#### 6.2 Trimming of MSAs at highly gapped positions

In order to test epistatic methods' sensitivity to noise arising from poorly aligned positions in the MSA, we analyzed ArDCA's performance on MSAs where highly gapped positions (columns) were removed. In all MSAs from dataset $\mathcal{D}$, we trimmed columns with a gap ratio above 0.2 and derived ArDCA predictions from the trimmed MSAs. This process removed approximately 13% of the positions in $\mathcal{D}$. Obviously, mutations in trimmed positions can no longer be predicted in this configuration, so we compared the performance of ArDCA on non-trimmed positions from both the full MSAs and the trimmed MSAs. As a result, we observed that using trimmed MSAs slightly improves performance, increasing the score from 0.474 to 0.484, while the scores of all independent-site methods, by definition, remain unchanged. Although the improvement is modest, it is noteworthy that removing data from the MSA can enhance performance. In other words, the predictive information provided by highly gapped positions is outweighed by the noise they introduce into the model. This observation supports the idea that the performance of epistatic methods in predicting protein stability changes is constrained by noise.

##### 6.3 Dependence on the regularization of epistatic contributions

Without going too deeply into the details of direct coupling analysis (DCA) methods, which are already well explained in the literature [13], let us give a brief overview of the main principles. DCA methods provide a statistical representation of a family of homologous protein sequences from a given MSA. Let $S = (a_1, a_2, \ldots, a_L)$ be a protein sequence of length $L$, where $a_i$ is the amino acid type at position $i$. We assume that the sequence $S$ is sampled across evolution with a probability given by the Boltzmann distribution, $P(S)$, which can be expressed as:

$$P(S) = \frac{1}{Z}\exp(-\beta\Phi(S)), \tag{1}$$

where $\beta$ is the inverse temperature, $Z$ denotes the partition function, and $\Phi(S)$ is the energy of the system:

$$\Phi(S) = -\sum_{1\leq i<j\leq L} J_{ij}(a_i, a_j) - \sum_{i=1}^{L} h_i(a_i). \tag{2}$$

In the previous equation:

- $h_i(a)$ are the independent-site field parameters, which intuitively define how well an amino acid type $a$ at position $i$ in the MSA fits within the DCA model.
- $J_{ij}(a_i, a_j)$ are the epistatic couplings and describe the likelihood of observing a pair of amino acids $a_i$ and $a_j$ at positions $i$ and $j$, respectively, in the MSA.

To infer the probability distribution and obtain the parameter values, maximum entropy approaches are used, ensuring that the probability of each residue at each position, as well as the joint probabilities of residue pairs at two positions, matches the observed frequency data in the MSA. As the inference of the full model is computationally demanding, some approximations are made. For example, in EVcouplings and pycofitness, the fields and couplings are estimated not from the likelihood of the probability distribution but by minimizing the negative log-pseudolikelihood $\mathcal{L}(h, J)$:

$$\mathcal{L} = -\log\left(\prod_{n=1}^{N_{\text{tot}}}\prod_{i=1}^{L} P(a_i|a_1..a_{i-1}, a_{i+1}..a_L)^{(n)}\right). \tag{3}$$

This is the product over the whole MSA of the conditional probabilities of observing residue $a$ at position $i$ given all other positions fixed. As there are often undersampling problems due to the number of parameters being large compared to the number of sequences in the MSA, regularization terms are usually added to the optimization function:

$$\mathcal{L}_{reg} = \mathcal{L} + \lambda_h \sum_{i=1}^{L}\sum_{a=1}^{21} h_i(a)^2 + \lambda_J \sum_{i=1}^{L}\sum_{j=1}^{L}\sum_{a=1}^{21}\sum_{b=1}^{21} J_{ij}(a,b)^2. \tag{4}$$

Typically, a constant regularization factor $\lambda_h$ is used for the $h$ coefficients, while a sequence-length dependent regularization factor $\lambda_J \propto (L-1)$ is used for the $J$ coefficients [6, 14]. Note that as the regularization parameter $\lambda_h$ (or $\lambda_J$) increases, the corresponding $h$ (or $J$) coefficients decrease. Once the model is inferred and all these coefficients are determined, the effect $\Delta X(i, a, b)$ of a single mutation at position $i$ from amino acid type $a$ to $b$ on the protein "fitness" is computed as:

$$\begin{aligned}\Delta X(i,a,b) &= \Phi(a_1,\ldots,a_{i-1},b,a_{i+1},\ldots,a_L) - \Phi(a_1,\ldots,a_{i-1},a,a_{i+1},\ldots,a_L) \\ &= [h_i(b) - h_i(a)] + \sum_{j=1}^{L} [J_{ij}(b,a_j) - J_{ij}(a,a_j)]\,. \end{aligned} \tag{5}$$

Using this equation, we can decompose the contribution of the DCA model into its independent-site contribution (denoted $\Delta h$) and epistatic contribution (denoted $\Delta J$). Additionally, the epistatic contribution can be further decomposed into the interactions between the mutated site and each other position $j$ in the MSA.

We performed several analyses with the regularization parameters using the pycofitness method. By fixing $\lambda_h$ to 10 and testing a range of different $\alpha_J = \lambda_J/(L-1)$ values between 0.2 and 50, we observed that the choice of regularization parameters can substantially impact the performance of pycofitness. It is interesting to note that $\alpha_J$ values between 6 and 50 result in relatively stable and optimal correlation values, which slightly outperform the independent-site $\text{LOR}_w$ score, with the optimal value being $\alpha_J = 15$ (Fig. S5a).

Interestingly, we found that different regularization strengths have opposite effects on predicting mutations in the core (RSA $<$ 20%) and on the surface (RSA $>$ 50%) of proteins (Fig. S5b). Weak regularization tends to improve core residue predictions, while strong regularization is optimal for surface residues. This result can be explained by the fact that core residues, having more interactions than surface residues, display a larger prevalence of epistatic effects and are thus better predicted by weak regularization (i.e., a higher contribution of $J$). The choice of regularization parameters is therefore not straightforward and can depend on the specific needs of the analysis.

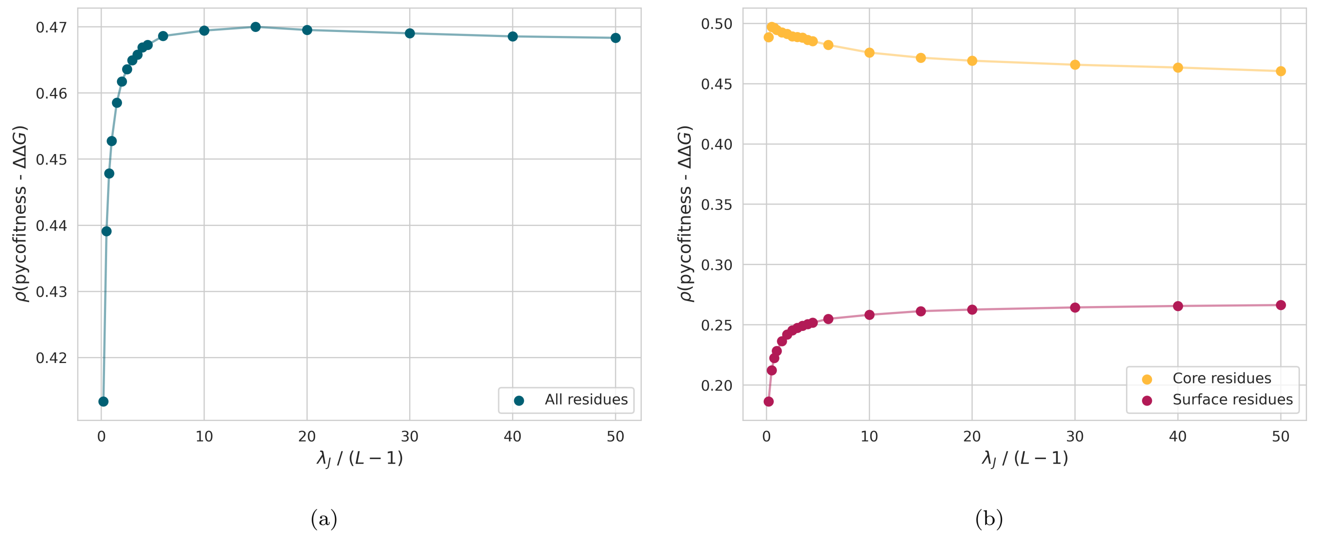

Figure S5: Average per-protein Spearman correlation coefficients $\rho$ between experimental $\Delta\Delta G$ values and pycofitness scores with regularization parameters $\lambda_h = 11$ and $\lambda_J$ ranging from 0.2 to 50 for: (a) all mutations from $\mathcal{D}$ and (b) core mutations (RSA $< 20\%$) in yellow and surface mutations (RSA $> 50\%$) in pink.

#### 6.4 Noise reduction in the coupling terms

Coupling values $J_{ij}(a, b)$ derived from DCA models provide valuable information about residue-residue interactions. However, the complexity of the DCA inference problem, which is generally based on a limited amount of sequence data relative to the number of $J_{ij}(a, b)$ parameters to optimize, makes these parameters noisy. While contact prediction primarily focuses on the top coevolving contacts, inferring the protein mutational landscape requires access to all couplings.

Analyses in the literature related to fitness prediction suggest that selecting a subset of couplings could improve method performance [9]. In this section, we show that reducing noise in the epistatic contributions by selecting only couplings corresponding to highly coevolving residue pairs improves the performance of DCA methods in predicting protein stability changes. As in the previous subsection, we applied this analysis to the pycofitness method.

The strength of the coupling between two positions $i$ and $j$ can be quantified by the Frobenius norm:

$$F_{ij} = \sqrt{\sum_{a,b} J_{ij}(a, b)^2}. \tag{6}$$

Using this norm, we modified the expression of coevolutionary energy $\Delta X_t(i, a, b)$ given in Eq. 5 by canceling the coupling terms if their Frobenius norm falls below a specified threshold $t$:

$$\Delta X_t(i, a, b) = [h_i(b) - h_i(a)] + \sum_{j=1}^{L} \delta(F_{ij} > t) \left[J_{ij}(b, a_j) - J_{ij}(a, a_j)\right], \tag{7}$$

where $\delta(F_{ij} > t)$ is equal to 1 if $F_{ij} > t$, and 0 otherwise.

We observe that removing weakly correlated couplings from the model enhances pycofitness performances on $\mathcal{D}$, improving the correlation from 0.469 to 0.484 for a threshold $t = 0.134$ (see Fig 5 in the main text). Although the increase in correlation is moderate, it is interesting to note that removing noisy couplings leads the epistatic models to outperform the independent-site model results (see Fig. 5 in the main text).

Furthermore, we observe that the optimal threshold value tends to increase with the $N_{\text{eff}}$ value of the MSA (Fig. S6). Smaller MSAs, being more prone to noise, exhibit an optimal threshold $t$ where most couplings are removed; conversely, larger MSAs have an optimal threshold $t$ where most couplings are retained. This further supports the idea that the performance of DCA-based methods in predicting protein stability changes is limited by noise.

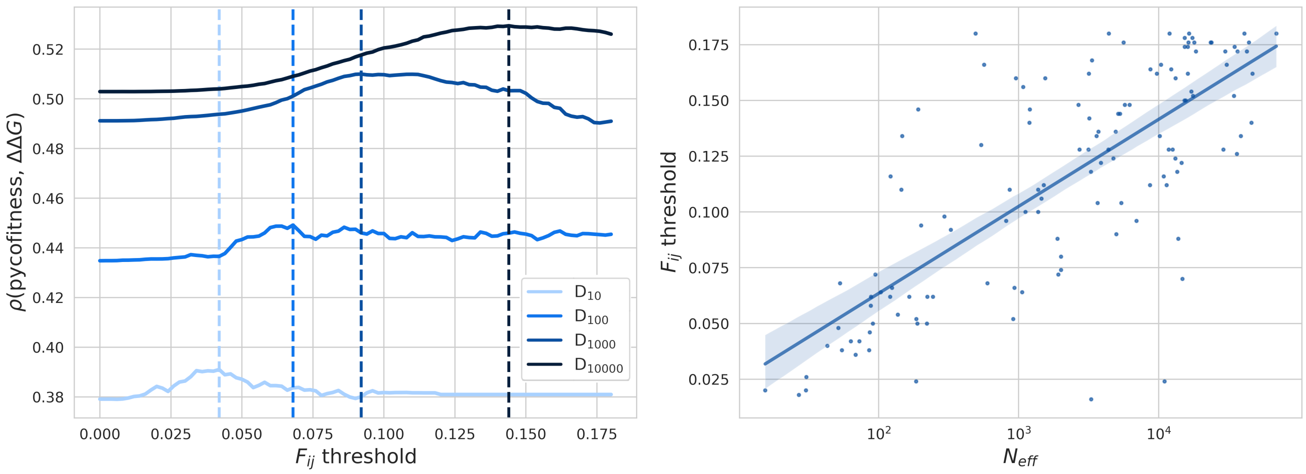

(a) (b)

Figure S6: Optimal Frobenius norm threshold $t$ dependence on $N_{\text{eff}}$. (a) Average per-protein Spearman correlation coefficient $\rho$ between experimental $\Delta\Delta G$ values and Frobenius norm-modified pycofitness score as a function of the threshold $t$ on the subsets $\mathcal{D}_{10}$, $\mathcal{D}_{100}$, $\mathcal{D}_{1000}$ and $\mathcal{D}_{10000}$. Vertical lines represent the optimal Frobenius norm threshold $t$ for each subset. (b) Per-protein optimal Frobenius norm threshold $t$ as a function of $N_{\text{eff}}$ (in $\log_{10}$ scale).

### 7 Performances of evolutionary models to predict $\Delta\Delta G$ on $\mathcal{D}$ subsets

We show in Tab. S4 the performance of different methods on subsets of $\mathcal{D}$ that depend on MSA depth. While the performance of evolutionary scores, alone or modulated by RSA, consistently increases with MSA depth, the performance of supervised $\Delta\Delta G$ predictors appears to be independent of the MSA depth (except for PremPS, which explicitly uses MSAs).

Table S4: Average per-protein Spearman correlation coefficients $\rho$ between experimental $\Delta\Delta G$ values and evolutionary scores or $\Delta\Delta G$ predictors' scores on the dataset $\mathcal{D}$ and its $N_{\text{eff}}$-dependent subsets $\mathcal{D}_{10}$, $\mathcal{D}_{100}$, $\mathcal{D}_{1000}$ and $\mathcal{D}_{10000}$.

| Method | $\mathcal{D}$ | $\mathcal{D}_{10}$ | $\mathcal{D}_{100}$ | $\mathcal{D}_{1000}$ | $\mathcal{D}_{10000}$ |
|---|---|---|---|---|---|
| GEMME-ind | 0.48 | 0.38 | 0.44 | 0.51 | 0.53 |
| pycofitness | 0.47 | 0.38 | 0.43 | 0.47 | 0.50 |
| $\text{LOR}_w$ | 0.47 | 0.37 | 0.44 | 0.48 | 0.50 |
| GEMME-epi | 0.46 | 0.37 | 0.42 | 0.48 | 0.49 |
| ArDCA | 0.45 | 0.25 | 0.39 | 0.50 | 0.53 |
| LOR | 0.45 | 0.35 | 0.43 | 0.47 | 0.50 |
| EVcouplings-epi | 0.43 | 0.36 | 0.42 | 0.43 | 0.47 |
| EVcouplings-ind | 0.42 | 0.36 | 0.42 | 0.42 | 0.45 |
| RSA $\odot$ GEMME-ind | 0.58 | 0.50 | 0.55 | 0.59 | 0.61 |
| RSA $\odot$ pycofitness | 0.59 | 0.50 | 0.56 | 0.61 | 0.63 |
| RSA $\odot$ $\text{LOR}_w$ | 0.59 | 0.51 | 0.57 | 0.60 | 0.61 |
| RSA $\odot$ GEMME-epi | 0.56 | 0.50 | 0.54 | 0.57 | 0.58 |
| RSA $\odot$ ArDCA | 0.60 | 0.50 | 0.57 | 0.62 | 0.64 |
| RSA $\odot$ LOR | 0.58 | 0.52 | 0.56 | 0.59 | 0.61 |
| RSA $\odot$ EVcouplings-epi | 0.59 | 0.51 | 0.57 | 0.61 | 0.63 |
| RSA $\odot$ EVcouplings-ind | 0.58 | 0.51 | 0.56 | 0.59 | 0.60 |
| PoPMuSiC | 0.61 | 0.62 | 0.64 | 0.59 | 0.60 |
| RaSP | 0.59 | 0.60 | 0.62 | 0.56 | 0.62 |
| PremPS | 0.59 | 0.54 | 0.58 | 0.57 | 0.63 |
| DDMut | 0.57 | 0.56 | 0.59 | 0.56 | 0.58 |
| KORPM | 0.55 | 0.58 | 0.55 | 0.52 | 0.55 |
| MAESTRO | 0.54 | 0.52 | 0.57 | 0.51 | 0.55 |
| DDGun3D | 0.53 | 0.55 | 0.53 | 0.52 | 0.52 |
